# Tropical montane and temperate hummingbirds incubate in small installments over long periods

**DOI:** 10.1101/2025.06.19.660382

**Authors:** Justin W. Baldwin, Mario Agustín Loaiza Muñoz, Juliana Sandoval-H, Scott K. Robinson, Ari Martinez, Gustavo Londoño

## Abstract

Life-history strategies and development are shaped by climate and predation. We examined how temperature and nest survival drive incubation behavior in 104 nests of 34 hummingbird species across 2,939m of elevation in the Peruvian and Colombian Andes. At high elevations, cold temperatures led to more frequent, brief incubation bouts and extended incubation periods; survival modulated incubation behavior with no effect on development. Hummingbirds had the highest egg surface-area to volume ratios among all birds; in 147 species, incubation periods increased with decreasing ambient temperatures. We propose a new hypothesis to explain how in cold conditions, heat transfer to the embryo is fragmented into smaller increments and therefore extended over longer periods, underscoring the role of adult behavior in mediating embryonic temperature mismatches and life-history tradeoffs.

## Introduction

Why some animals reach maturity in days while others take years is a fundamental question in ecology and evolution (*1, 2*). Developmental duration, a key dimension of life history (*3*), is shaped by mechanistic and physiological constraints (*4*–*6*), phylogeny (*7*), environmental conditions and survival rates (*8, 9*). In birds, the incubation period is an energetically expensive phase of development (*10, 11*) during which adults balance competing demands. Adults must ensure that egg temperatures remain high and stable (∼34-40.5° C, well above “physiological zero” of 26° C (*12*–*15*)). Species with uniparental care, however, must leave the nest to refuel; during this time, developing embryos are subjected to cold and variable temperatures and incur fitness costs (*16*). Harsh conditions, rain and food scarcity can extend parental absences and decrease egg temperatures (*15, 17*) factors that slows development (*16, 18*). Exacerbating these challenges, the threat of nest predation forces adults to conceal (*19*) or limit nest visits (*20, 21*), and selects for accelerated development (*3, 4*). When predation pressure is relaxed, incubation behavior changes (*22*) and development is prolonged (*23, 24*). Nevertheless, the relative importance of abiotic conditions and nest survival rates on incubation and life history remains uncertain (*9*).

Our understanding of the tradeoff between abiotic and biotic constraints on incubation results is limited for several reasons. Tropical birds are widely thought to have uniformly low rates of nest attendance (*17, 18, 25*), but incubation behavior from many tropical species remains unknown (*26*). Additionally, temperate-tropical comparisons often use nest attentiveness, the fraction of daytime that birds spend incubating on the nest (*12, 25*), but this metric alone might be insufficient: diurnal incubation can be fragmented into many short or few long bouts, maintaining nest attentiveness (Fig. S1) but also likely impacting egg temperatures (*16*) and predation risk (*20*).

Montane hummingbirds in the Neotropics offer a unique opportunity to disentangle the joint effects of environmental conditions and nest predation on incubation behavior and life-history traits. Hummingbirds are extremely sensitive to changes in environmental temperatures because powered hovering flight consumes large amounts of energy (*27, 28*), and much of the resulting heat produced is lost to the immediate environment due to the birds’ small size and high ratio of surface-area to volume (*29, 30*). The different temperature regimes they encounter as they breed from sea-level to above 4,000m may affect their range of body size (1.9–20.2g), incubation period (12–23 days) and egg mass (0.25–1.75g) (*6, 31*–*34*). Moreover, some major traits that explain life-history variation across birds (*34*) are controlled for in hummingbirds: all are nectarivorous and have 2-egg clutches, uniparental care and similar nest architecture (*34*). Embryonic development might be especially sensitive to environmental temperatures and highly dependent on parental incubation, because their small eggs rapidly lose heat when unattended (*35*).

In this study we examined how nest survival rates and environmental temperatures influence the incubation and development of 34 species of Neotropical hummingbirds, using observations from 293 complete days of incubation in104 nests, spanning 2,939 m of elevation in the Peruvian and Colombian Andes. We hypothesize that hummingbirds are uniquely temperature-sensitive due to high ratios of egg surface-area to volume and therefore employ locally adaptive incubation strategies across the elevational gradient. Accordingly, hummingbirds at cold, high elevations should employ more frequent, brief incubation bouts compared to species at warm, low elevations (*9, 15, 36*). Moreover, long incubation recesses, which expose embryos to cold temperatures that slow development (*16, 37*), are likely compensated for by extended incubation periods and increased egg mass, following the “embryonic temperature hypothesis” (*18*). Additionally, we explored how elevational variation in nest survival rates (*38*) shapes both incubation behavior (*20*) and incubation periods using estimates of nest survival from 390 nests of the same 34 species. We test these predictions with comparisons of incubation behavior, incubation periods, egg mass and egg shape using our new data from the field and data from the literature (*6, 39*–*41*).

## Results and Discussion

### Hummingbird incubation behavior in the Andes

We used in situ dataloggers and thermocouples inserted into active nests (Fig. 1A) to determine incubation behavior from 104 nests of 34 species of hummingbirds during eight years of fieldwork at eight field sites in Peru and Colombia (Fig. 1B-D) (*42*). We obtained a mean of 2.82 full days of incubation per nest (SD = 1.46, range 1-16 days, 293 days total). Incubation behavior varied within and across species and along the elevational gradient (Fig. 1E): in high-elevation species (e.g., Metallura tyrianthina) incubation bouts were frequent (mean = 82.6 trips/day, range: 49-183) and brief (mean = 11.3 min, range: 4.8-14.7), but in comparison, in low-elevation species (e.g. Glaucis hirsutus) incubation bouts were fewer (mean = 25.1 trips/day, range: 15-39) and extended (mean = 40.3 min, range: 11.6-69.3).

**Fig. 1.**
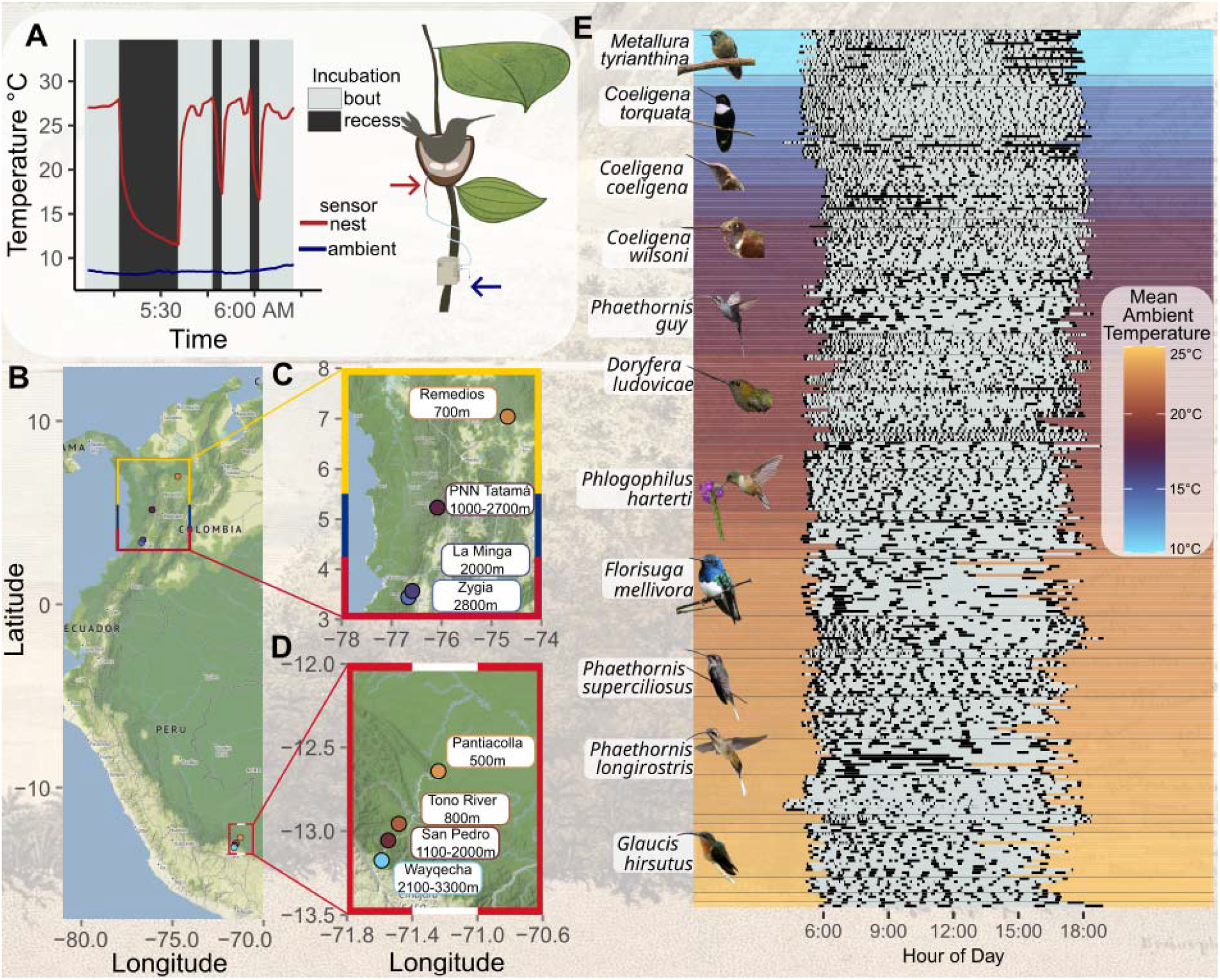
Incubation behavior of montane hummingbirds in Colombia and Peru along an elevational gradient. (**A**) Active nests were monitored using remotely deployed data loggers with thermocouples inserted below eggs to collect nest (red) and ambient temperatures (blue). Temperature traces revealed incubation bouts (grey) and recesses (black; here an Amethyst-throated Sunangel, *Heliangelus amethysticollis*). (**B**) Eight years of fieldwork across 2,939 meters of elevation in the Andes spread across four field sites in Colombia (**C**) and Peru each (**D** yielded 293 full days of incubation behavior from 104 nests of 34 species (**E**). Horizontal thin grey lines separate days from same species. Grey rectangles show incubation bouts and black rectangles show incubation recesses. (E) Days of incubation are ordered by species and daily ambient temperature, with station (B-D) mean diurnal ambient temperatures visualized according to the same color scale. Photos of well-sampled species provided by Lina Peña Ramirez, Carlos Calle and contributors to Wikimedia Commons (see acknowledgments) under Creative Commons licenses. Icon (A) made by Isabella Burgos. Basemaps (B-D) from Stadia/OpenStreetmap.

### Ambient temperatures shape hummingbird incubation behavior and life history

Hummingbird eggs likely lose heat faster than other species, due to their small size and uniquely cylindrical (“tic-tac”) shape. We predicted that the ratio of egg surface-area to volume is higher in hummingbirds than in other birds. In a comprehensive sample of egg dimensions (*41*), hummingbirds had the highest ratio of egg surface-area to volume among birds worldwide (Trochilidae; mean = 2.27, SE = 0.025, N = 12): 3.8 times higher than the average family (thrushes, Turdidae; mean = 0.59, SE = 0.03, N = 36), and ∼89 times higher than the family with the lowest surface-area to volume ratio (ostriches: Struthionidae; mean = 0.026, N = 1; Fig. 2A). Accordingly, we posit that hummingbird eggs are highly temperature-sensitive and dependent on adult incubation behavior.

**Fig. 2.**
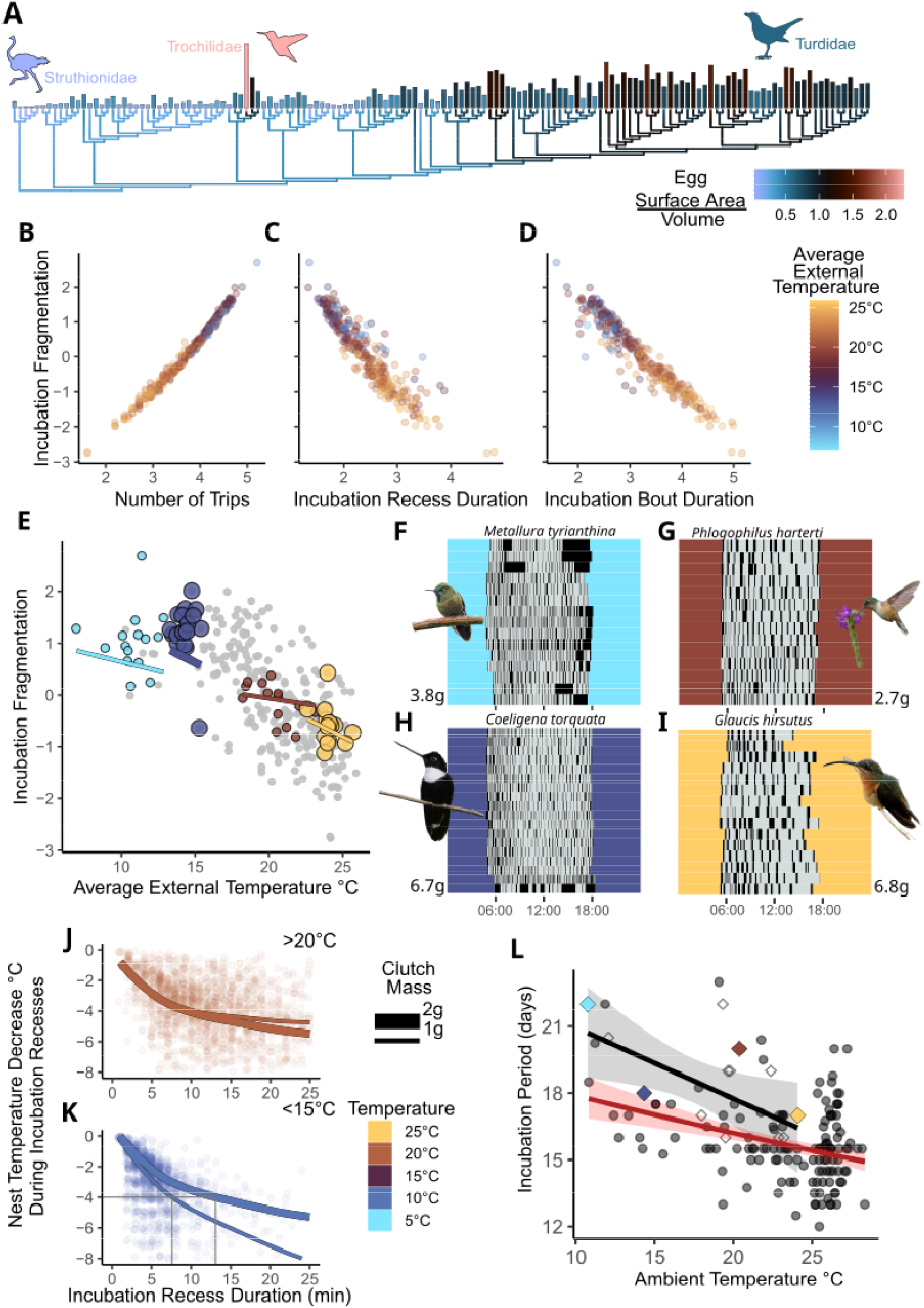
Thermally sensitive hummingbird eggs demand temperature-dependent incubation strategies; these in turn shape development. (**A)** Hummingbirds have the largest ratios of egg surface-area to volume across all bird families (Trochilidae mean = 2.27, SE = 0.025, N = 12). Icons indicate mean egg surface-area to volume for ostriches (mean = 0.0256) and thrushes (mean = 0.593, SE = 0.003, N = 36). High levels of incubation fragmentation are characterized by many incubation trips (**B**), brief incubation bouts (**C**) and brief incubation recesses (**D**) – all traits on x-axis were natural-log transformed. Phylogenetic generalized linear mixed models reveals that temperature affects incubation fragmentation and interacts with body mass (**E**). Small-bodied (**FG**) and large-bodied species (**HI**) are shown from high and low elevations, respectively. Decreases in nest temperature were driven by nonlinear effects of incubation recess duration as well as interactions with ambient temperature and clutch mass (**JK**). Small eggs in cold temperatures experienced the largest decrease in temperature, dropping 4 degrees in 7.6 minutes, whereas larger eggs dropped 4 degrees in 13.1 minutes to (thin grey lines - K). (**L**) Phylogenetic regressions show prolonged incubation periods in cold temperatures in 147 species worldwide (estimate = -0.041, SE = 0.010, T = -4.303, p< 0.001, red line) and in 13 species on the elevational gradient (estimate = -1.234, SE = 0.527, t = -2.323, p = 0.039, black line, colored diamonds correspond to species in F-I). We thank Felix Uribe and Charles J. Sharp for providing photos on Wikimedia Commons under Creative Commons licenses as well as Carlos Calle for the photo of *Phlogophilus*. Silhouettes in A from PhyloPic and we thank Charles J. Sharp for providing photos on Wikimedia Commons under Creative Commons licenses.

Across the elevational gradient, the daily number of incubation bouts and recesses, as well as their duration, were interdependent, so we used principal component analyses to define an aggregate metric, hereafter “incubation fragmentation” (Table S1). This approach allowed us to separate days of incubation behavior with few long incubation bouts from days of incubation behavior fragmented into many, brief bouts (Fig. 2B-D). Phylogenetic generalized linear mixed models of incubation fragmentation revealed a strong significant effect of ambient temperature (posterior mean = -0.486, 95% credible interval: -0.657 – -0.314, Table S2, Fig. 2E). But when temperatures rose, incubation fragmentation decreased more in large species than in small species (temperature x mean body mass interaction, posterior mean = -0.166, 95% credible interval: -0.322 – -0.006, Table S2, Fig. 2F-I). Incubation fragmentation was unrelated to traditional measures of nest attentiveness (Pearson’s correlation coefficient = -0.015, t = -0.254, df = 275, p-value = 0.800, Fig. S2), and analysis of nest attentiveness (instead of incubation fragmentation) revealed no influence of temperature (Table S3). Our results were robust to including phylogenetic relatedness in the construction of principal components (Table S4) and phylogenetic uncertainty (Table S5).

Although during incubation recesses, nest temperatures eventually approximated ambient temperatures (Fig. S3), nests with small eggs lost temperature faster – but only when ambient temperatures were cold (three-way interaction between non-linear effect of incubation recess duration, ambient temperature and clutch mass: F_5737,1_ = 81.902, p-value < 0.0001, Table S6). During warm ambient temperatures, clutch mass conferred no thermal advantage (Fig. 2J). In contrast, during cold temperatures (10°C), large clutch mass buffered temperature loss: for example, a species with 2g clutch mass lost 4°C of nest temperature during a 13.1-minute incubation recess, whereas a species with 1g clutch mass lost the same amount in 7.6 minutes (Fig. 2K). As body mass and egg mass scale allometrically (*43*), the interacting effects of temperature and body mass on incubation behavior (Fig. 2E) could be explained by the way large-bodied birds with correspondingly large eggs (these have low egg surface-area to volume ratios; Fig. 2A) can decrease incubation fragmentation (and extend the duration of their incubation recesses) without exposing their embryos to low temperatures. Our results were similar when accounting for phylogenetic and temporal non-independence, although the support for some parameters decreased (Table S7, Fig. S4).

Across the elevational gradient, development accelerated with increasing local ambient temperatures: the incubation period decreased from 22 days at 10.7°C (2,879m) to 16 days at 22.9°C (431m) (univariate phylogenetic regression: estimate = -1.234, SE = 0.527, t = -2.323, p = 0.039, N = 13; Fig. 2L diamonds, black line). Across the western hemisphere, hummingbirds in colder areas also had significantly longer incubation periods than those in warm areas (temperature estimate = -0.041, SE = 0.010, t = -4.303, p < 0.001, N = 147, Table 1, Fig. 2L black points, red line). Previous studies have found inconsistent relationships between nest attendance and incubation period (*12, 25*) and species with long incubation recesses have long incubation periods (*12, 18*). However, our results suggest that in cold ambient temperatures, incubation recesses are short whereas incubation periods are long. This is additionally surprising as temperature has no influence on incubation periods across all birds (*6*). We also recovered the expected allometry (*5, 6*) between incubation period and body mass (Table 1). Temperature had no effect on egg mass (estimate = -0.025, SE = 0.020, t = -1.212, p = 0.217, N = 104, Table S8). However, we found a weak, yet marginally significant positive relationship between egg mass and incubation period, after accounting for body mass (estimate = 0.112, SE = 0.060, t = 1.878, p = 0.064, N = 89, Table S9, Fig. S5), which suggests that increased parental investment in egg mass is reflected in extended incubation periods (*18*). This correlation is surprising, as egg mass and size are likely under strong physical constraints in hummingbirds (*44*). Our inferences were robust to phylogenetic uncertainty (Table S10).

**Table 1.** Temperature, body mass and insularity influence hummingbird incubation periods. Phylogenetic generalized linear models of incubation periods. Significant parameters in bold. ⍰ Denotes terms eliminated during model selection. Adjusted R^2^=0.235 and λ = 0.221 in final model.

| Parameter | N | Estimate | Standard Error | T-value | p-value |
| --- | --- | --- | --- | --- | --- |
| Nest Height □ | 112 | 0.009 | 0.011 | 0.810 | 0.420 |
| Habitat: low □ | 145 | -0.019 | 0.040 | -0.465 | 0.643 |
| Habitat: medium □ | 145 | -0.052 | 0.380 | -1.358 | 0.177 |
| Habitat: high □ | 145 | -0.034 | 0.399 | -0.860 | 0.391 |
| Migration □ | 145 | -0.012 | 0.023 | -0.534 | 0.594 |
| Precipitation □ | 147 | 0.001 | 0.010 | 0.132 | 0.895 |
| Latitude □ | 147 | -0.009 | 0.009 | -0.951 | 0.343 |
| **Insularity** | 147 | 0.088 | 0.030 | 2.899 | 0.0043 |
| **Body Mass** | 147 | 0.044 | 0.008 | 5.219 | <0.0001 |
| **Temperature** | 147 | -0.041 | 0.009 | 4.761 | <0.0001 |

### Nest survival shapes hummingbird incubation and life history

We additionally obtained estimates of nest survival rates from 390 nests of 34 species by exploring 6,092 cumulative days of nest observations; during this time we registered 178 instances of predation. When data were pooled across the gradient and within stations, aggregated Mayfield estimators (*45*) of daily survival rates were high (Fig. 3A, gradient: 0.957, SE = 0.003, N = 390 nests; range: Pantiacolla : 0.948, SE = 0.005, N = 170, Tatamá: 0.967, SE = 0.006, N = 47; Table S11) - higher than other cup-nesting species and near the highest values of cavity-nesters (*39*). Predation rates were low (Fig. 3B, Table S11). To correlate survival with incubation behavior, we modelled daily nest-level survival and predation rates using phylogenetic logistic exposure models (*46*). Shared ancestry (i.e., phylogenetically pooled random intercepts) explained substantial variation (Figs. 3CD, Fig. S12). In contrast, neither nest height (posterior mean = 0.07, 95% credible interval: -0.79 – 0.87), elevation (posterior mean = 0.03, 95% credible interval: -0.37 – 0.44) nor interannual variation significantly affected daily survival rates (Fig. 3E-G, Table S12). Daily predation rates were likewise unaffected by ecological variables (Table S13, Fig. S6). Daily survival rates had a significant positive effect on incubation fragmentation (posterior mean = 0.144, 95% credible interval: -0.266 – -0.019; Fig. 3H, Table S2), suggesting that when nest survival is low, visits to the nest were less frequent and lasted longer. These findings support a longstanding idea that predation risk influences incubation behavior (*20*). Our regression analyses were robust to the use of predation rates instead of survival rates (Table S14).

**Fig. 3.**
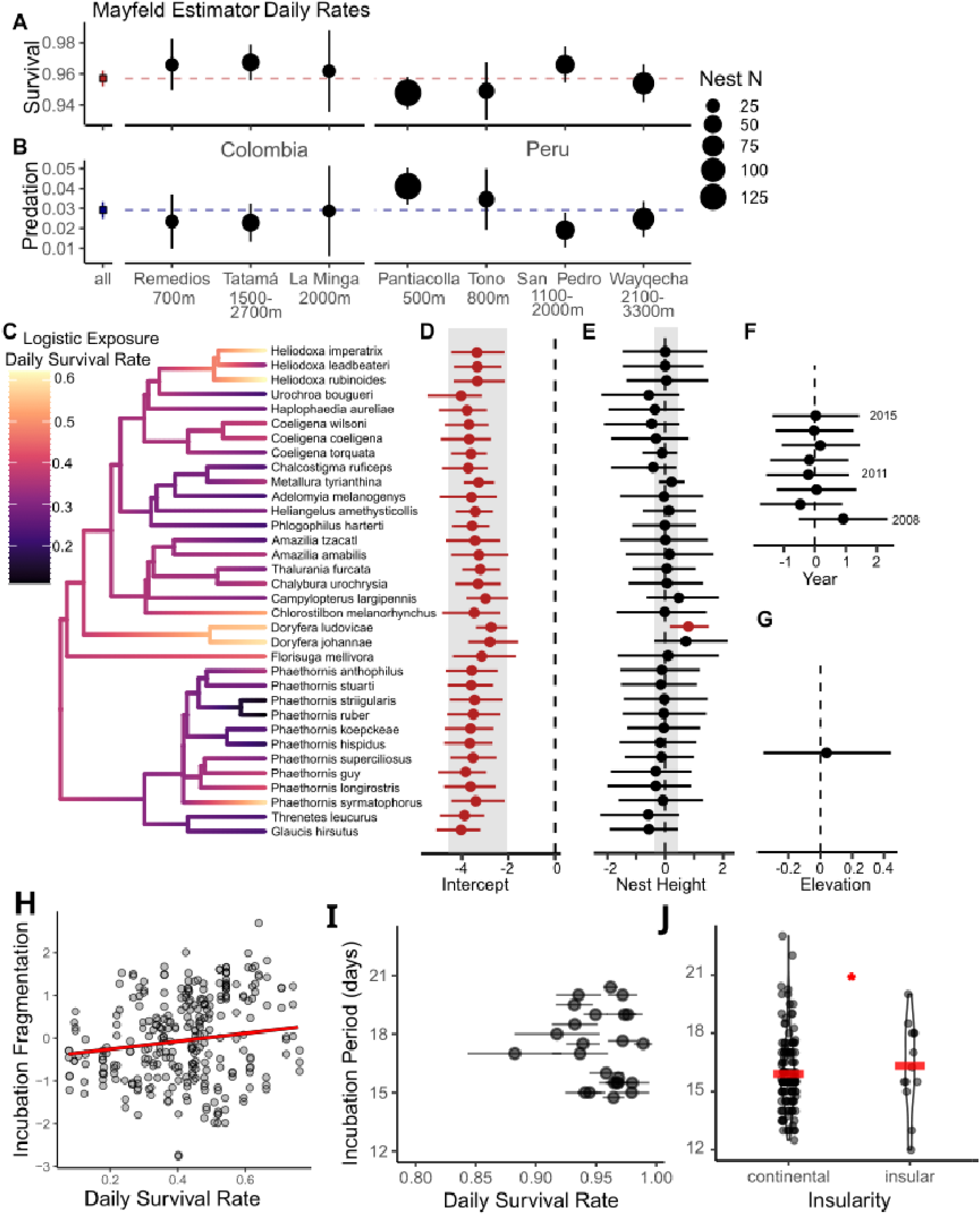
Nest survival rates of montane hummingbirds vary with phylogeny, and influence incubation behavior but not development. Mayfield estimators of daily survival (**A**) and predation rates (**B**) along the elevational gradient where N denotes number of nests. Horizontal dotted lines denote pooled estimates across gradient. (**C**) Phylogenetic logistic exposure models reveal notable variation in daily survival rates across species, with weak species-specific effects of nest height. Vertical dashed lines indicate zero. Tip color denotes species-average survival estimate of predicted values for each nest (C). (**D-G**) Points show species-specific posterior means, and horizontal lines show 95% credible intervals of effect sizes from phylogenetic logistic exposure models, with significant effect sizes in red. Shaded rectangles show hyperparameter distributions for random intercepts (D) and random slopes of nest height (E). Effects of interannual variation (F) and elevation (G) applied to all species equally. Across the elevational gradient, incubation fragmentation increased with increasing daily rates of nest survival (**H**). Survival rates had no effect on incubation periods (**I**, estimate = 0.009, SE = 0.029, T = 0.327, p = 0.750, N = 30, horizontal lines denote standard errors). Island-dwelling hummingbirds have significantly longer incubation periods than continental species (**J**, estimate = 0.097, SE = 0.031, T = 3.171, p = 0.0019, red lines show means).

Given that low survival caused birds to spend longer incubation bouts on the nest (keeping egg temperatures high), we predicted that eggs in such nests would develop faster, in line with life-history theory (*3, 23*), and because long incubation bouts could keep egg temperatures high (*18*). To test the influence of nest survival on incubation period, we performed a regression analysis on the 30 species of hummingbirds for which both daily survival rates and incubation periods are available (*39*). These results revealed no effect of survival rates on incubation periods (estimate = 0.009, SE = 0.029, t = 0.327, p = 0.750, N = 30, Table S15, Fig. 3I) and instead confirmed the significant effect of ambient temperature (Table S15). These conclusions were robust to measurement error in survival estimates (Table S16).

In addition, insular species had on average 1.1-day longer incubation periods than continental species (estimate = 0.097, SE = 0.031, t = 3.171, p = 0.002, N = 147, Table 1, Fig. 3J). Although this effect size is small, it corroborates the “slow pace of life” on islands (*6, 23, 47, 48*). Overall, while nest survival rate does influence incubation behavior in hummingbirds, its effect on their life-history traits seems weaker than that of ambient temperature, although more data are needed. This relatively small effect of variation in survival rate on nesting behavior contrasts with patterns in other tropical families (*21, 49*). We suspect that hummingbirds might be uniquely unaffected by nest predators because of their allometrically unparalleled flight speeds (*50*), pugnaciousness (*51*) and ability to nest in enemy-free areas (*52*). Additionally, predators prefer large eggs (*53*), which offer more protein: in context, a nest predator in the Peruvian highlands would consume the same egg mass from a single Andean Guan egg (Penelope montagnii, 73.4g, N = 1), as from ∼7 Great Thrush eggs (Turdus fuscater, mean = 10.5g, SE = 0.114, N = 86) or ∼146 Tyrian Metaltail eggs (Metallura tyrianthina, mean = 0.504g, SE = 0.014, N = 49; Fig. S7).

### The Incubation Fragmentation Hypothesis

Our results add a novel thermal perspective to hummingbird life history and reveal extreme, unexpectedly variable incubation behavior in the Neotropics in a single avian family. Along the elevational gradient, colder temperatures fragmented incubation behavior (Fig. 1E, Fig. 2B-I). Likewise, global temperature variation affected incubation fragmentation in a sample of 201 species from 37 families (*36*) (phylogenetic regression: estimate = -0.146, SE = 0.037, t-value = -3.950, p = 0.00001, λ = 0.989, N = 201; Fig. S8), suggesting a universal effect of temperature on incubation behavior. Our finding -- that across the globe hummingbirds in cold temperatures (where incubation recesses are frequent and brief) require extended incubation periods (Fig. 2DL, Table 1) – is then particularly surprising, as the “embryonic temperature hypothesis” (hereafter ETH) predicts that species with protracted incubation recesses should have extended incubation periods (Fig. 4A) due to cold exposure during extended parental absences (*12, 18*).

**Fig. 4.**
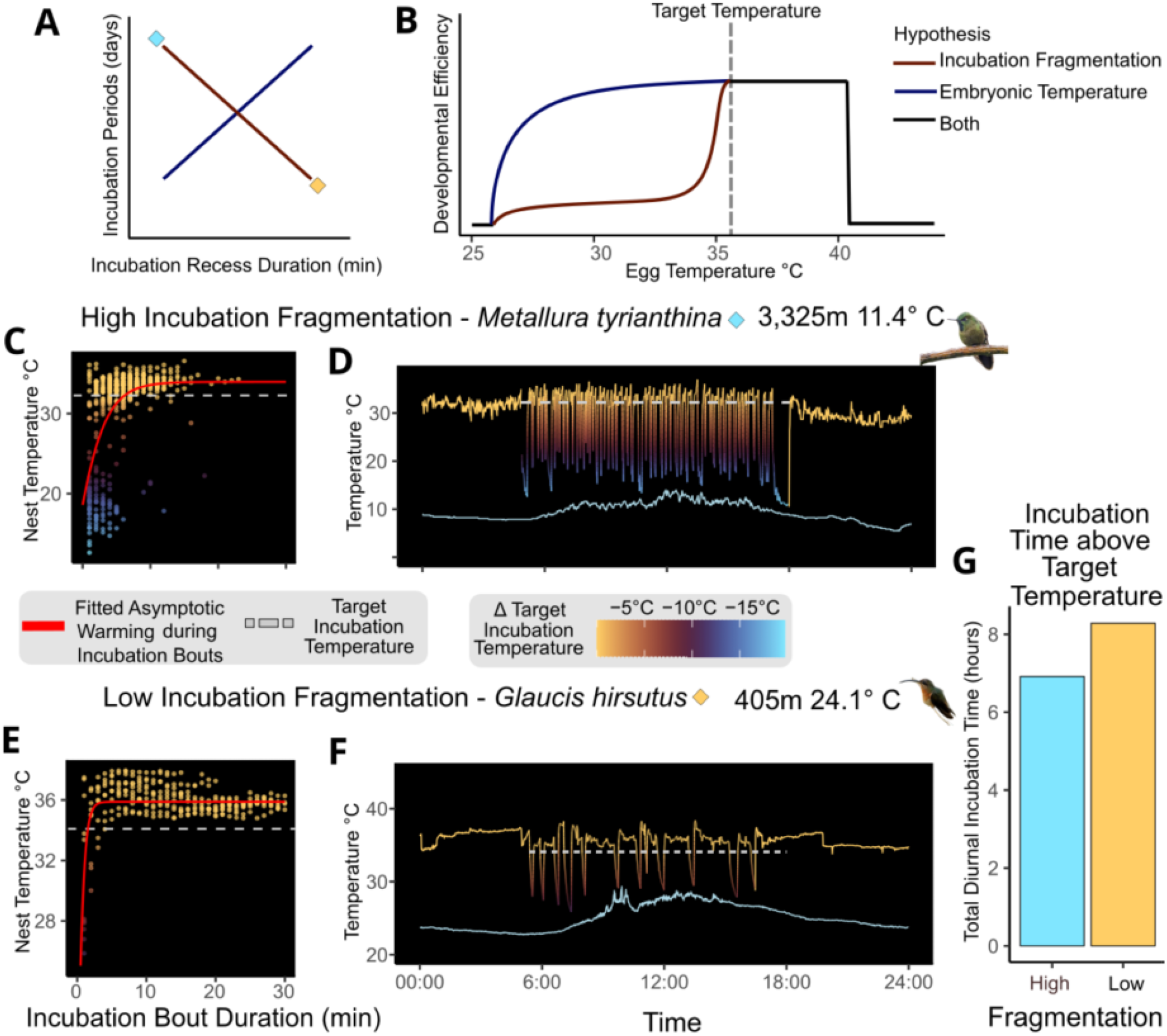
The Incubation Fragmentation Hypothesis. (**A**) The IFH and ETH make different predictions about the relationship between incubation recess duration and incubation period, possibly due to the differently-shaped relationships (**B**) between temperature and embryonic developmental efficiency. (B–ETH curve adapted from (*15*)). (**C-F**) Asymptotic warming during incubation bouts reveals nest temperatures approaching target temperatures for species with high and low incubation fragmentation (C,E). Target temperature (dashed grey line) is 5% below asymptote estimates from Gompertz functions (red lines) fitted to nest temperatures during diurnal incubation bouts. (D,F) Nest temperature traces are colored by distance from target temperatures; ambient temperatures shown in pale blue line. (**G**) Nest temperatures of species with high incubation fragmentation (blue diamond in A) are less frequently above target incubation temperatures on a daily basis compared to nest temperatures of species with low incubation fragmentation (yellow diamond in A). Species with high levels of incubation fragmentation take longer to complete the incubation period (A).

To explain our results, we propose an alternative hypothesis, the “incubation fragmentation hypothesis” (IFH). Under both hypotheses, endothermic adult birds provide close physical contact with eggs during incubation, rapidly transferring heat to their ectothermic embryos. Embryonic temperature influences growth efficiency (*54*), which is highest at incubation temperatures near adult body temperatures (34–40.5°C (*12–15*)). The hypotheses differ in how embryonic growth efficiency decreases when temperatures drop below this range (Fig. 4B). Citing poultry science (*14*), the ETH assumes convex decay of growth efficiency with decreasing egg temperature (*15, 18*). But as embryonic temperature thresholds and thermal sensitivity vary strongly across species (*13*), we propose that growth efficiency might decay asymptotically in hummingbirds (Fig. 4B). Even if hummingbirds also show convex decay, their high surface-area to volume ratios could rapidly drop embryo temperatures below target incubation temperatures soon after incubating adults leave the nest (Fig. 1A, Fig. 4CD, Fig. S9), particularly impacting small species in cold environments (Fig. 2JK). Additionally, frequent interruptions of incubation might stunt any developmentally essential processes that are initiated only after sustained exposure to target incubation temperatures (i.e. duration-dependent regulation (*55, 56*); Fig. S10).

Consequently, species with high (Fig. 4CD) and low rates of incubation fragmentation (Fig. 4EF) develop differently: frequent, short incubation recesses decrease the total time an embryo spends at developmentally relevant incubation temperatures (Fig. 4G, lower daily “cooking time” (*16, 57*)); extended incubation periods compensate for the lower daily thermal dose (Fig. 4A, 2EL) and increased egg mass (Fig. S5) as in (*18*). Whereas the ETH highlights reduced embryonic growth during incubation recesses, the IFH emphasizes increased embryonic growth during long incubation bouts. However, as number and duration of incubation recess and of incubation bouts are interdependent (Fig. 2B-D), this difference in emphasis produces conflicting predictions about the effect of behavior on development (Fig. 4A).

Although we were unable to formally test the ETH without hummingbird egg temperature measurements across all elevations, we evaluated two corollary predictions. Provided ambient temperatures are well below egg temperatures (*18*), long incubation recesses alone can reduce egg temperatures and delay development (*12*), supporting the ETH. We could not readily measure hummingbird egg temperatures due to their size, so we used a limited sample of two nests. All nest and egg temperatures in these nests were significantly correlated (all ρ > 0.6, p < 0.001; Fig. S11), and target egg temperatures exceeded 28° – well above local mean ambient diurnal temperatures (< 20.2° C). In other words, the predictions of the ETH were expected (Fig. 4A). Moreover, variation in target egg temperature across elevations might explain differences in incubation period (*12*), and body temperature limits target egg temperature. Yet in over 877 hours of internal body temperatures from 112 non-torpid hummingbirds (24 species, from 520-2,500m in Colombia (*58*)), nocturnal body temperatures did not vary across elevation (mean = 37.07° C, range = 30.02-41.05; phylogenetic mixed model: elevation effect = 0.21, SE = 0.52, 95% CI = 0.10-2.08, Table S17, Fig. S12). Therefore, we infer that target egg temperatures during incubation neither vary across elevation nor explain variation in incubation periods in hummingbirds.

Avian life-history research remains an active area of research; our results suggest that our understanding of these processes will improve greatly as we acquire more data using new technologies for poorly known tropical species (*26*). With our new data on the duration and frequency of incubation bouts, we can now understand how parental behavior affects multiple life-history tradeoffs in response to both abiotic (temperature) and biotic (predation) conditions (*59*). Mismatches between optimal and realized developmental temperatures during incubation can incur myriad costs such as increased mortality, decreased developmental speed (*60*), reduced immunocompetence (*61*), altered locomotor performance (*62*) and increased fearfulness (*63*). As global temperatures continue to change, such mismatches might become more frequent and could influence species survival and population persistence. Overall, our results speak to wide-ranging effects of avian parental behavior that have the potential to shape evolutionary processes via life-history tradeoffs, particularly when behavioral strategies buffer organisms from their environment (*64, 65*).

## Acknowledgments

We thank everyone who supported these experiments; see supplementary materials for a full list.

## Funding

National Science Foundation grant DEB-1120682 (GAL)

Association of Field Ornithologists, Alexander Skutch Award (GAL)

Wilson Ornithological Society, Louis Agassiz Fuertes Award (GAL)

American Ornithologists’ Union, Alexander Wetmore Award (GAL)

Fulbright, Student Research Scholarship (JWB)

## Author contributions

Writing – review & editing: JWB, SKR, AM, GAL

## Competing interests

Authors declare that they have no competing interests.

### Data and materials availability

All data and code are available on Zenodo (DOI: 10.5281/zenodo.15670484, https://zenodo.org/records/15670485) and detailed methods are in the Supplemental Materials.

## Supplementary Materials

Materials and Methods

Supplementary Text

Figs. S1 to S12

Tables S1 to S17

References (*66*–*102*)

## Notes

https://zenodo.org/records/15670485

### Competing Interest Statement

The authors have declared no competing interest.

