## Supplementary material for "Tropical montane and temperate hummingbirds incubate in small installments over long periods": Methods and Supplemental Material

**This PDF file includes:**

Materials and Methods

Figs. S1 to S12

Tables S1 to S17

References (*66-102*)

***1. Data and data sources***

All data and code are available on Zenodo (DOI: 10.5281/zenodo.15670484, https://zenodo.org/records/15670485).

***1.1. Field sites.*** Nesting data were collected during eight years of fieldwork between 2008 and 2015 in Peru and Colombia. Between 2008 and 2014 we worked at four field sites in southeast Peru, which were distributed along the elevational gradient of the Kcosñipata Valley on the east slope of the Andes (Fig. 1). Each year, from August to December, we visited the following sites with teams of nest-searchers: Pantiacolla Lodge (412 m asl), the Tono River (950 – 1,000 m asl), the Cock-of-the-Rock Lodge in San Pedro (1,100 – 2,000 m asl), and Wayqecha Research Station (2,300 – 3,100 m asl). All study sites are connected by continuous forest and form an intact elevational gradient. For additional details on the Peruvian study sites, see (*42*, *66*). Between 2014 and 2015, we also worked at four field sites in west and central Colombia. Teams of nest-searchers deployed variably throughout the year to the following stations: Remedios (700 m asl, Antioquia), Montezuma Rainforest Lodge in Tatamá National Park (1,100 – 2,700 m asl, Risaralda), La Minga Ecolodge (2,000 m asl, Valle del Cauca) and Zygia Icesi University Field Station (2,300 – 2,800 m asl, Valle del Cauca). For additional information on Colombian sites, see (*58*, *67*–*69*).

***1.2. Nest searching and monitoring.*** At each study site, field crews of ca. 5 technicians searched for nests for six days per week, ten hours per day. Each technician was assigned a unique 10-to-15-hectare plot where they would look for nests during the season. When a nest was discovered, nest monitoring would commence using either camera traps, visual monitoring or temperature loggers (next section), depending on what equipment was immediately available.

Professional grade infra-red sensitive color camera traps (Reconyx PC85 or R60 Rapidfire models, Reconyx, WI, USA, <https://www.reconyx.com>) were installed ca. 50 – 100 cm away from the nest on tripods or suitable branches and camouflaged with nearby vegetation. We programmed camera traps to shoot 10 photos upon motion detection at the nest to capture fast-moving predators and also programmed cameras to take 1 shot per minute to detect slow-moving reptilian predators. Cameras remained deployed until nest failure or completion. Furthermore, the presence of camera traps at the nest and variation in monitoring techniques had no effect on nest predation rates (*42*). We inspected nests visually with in-person visits every three days during incubation phase and every day or every other day during the nestling phase. Nest visits concluded when all nestlings fledged or nest contents were depredated.

Each nest could terminate in one of three events: success, nest failure, or predation. Successful nests had active and fully feathered nestlings that eventually disappeared between visits without evidence of predation. Depredated nests were ones where eggs or nestlings disappeared at premature developmental stages with evidence of predation events and/or visual corroboration of the predation event from the camera traps. Failed nests were ones where either eggs did not hatch or nestlings died for other reasons (e.g. water levels rising on river, extreme precipitation and wind knocking down nest, etc.). We readily acknowledge that our monitoring techniques could bias estimates of survival, but as our protocol was applied to all sites and years equally, we expect any bias to be non-systematically distributed.

***1.3. Quantification of incubation rhythms with temperature loggers.*** Our third method for following nest activity and identifying nest fate also generated data on incubation behavior. We used thermocouples connected to U-12 HOBO data loggers (Onset Computer Corporation, Pocasset, MA, USA, <https://www.onsetcomp.com>), which we programmed to take one temperature measurement per minute. Each data logger was connected to two thermocouples, one of which was placed inside the nest under the eggs and the other of which was placed on the exterior of each nest (Fig. 1A). This allowed us to identify the incubation on-bouts by sharp temperature increases that plateaued and incubation recesses by sharp temperature decreases that eventually approximated the ambient temperature (Fig. 1A). We then used Raven Pro software ((*70*) https://www.ravensoundsoftware.com/) to process the temperature traces and generate incubation rhythms. For more details on data processing, see (*71*). Three of us (MALM, JSH, JWB) visually cross-checked the raw temperature with the incubation rhythms to ensure data quality. We also developed a custom R Shiny Application to manually process data streams and generate incubation rhythms when the incubation rhythms generated by Raven Pro failed our data quality check. As incubation behavior can change slightly throughout the incubation period, we selected, when possible, up to three complete days of incubation behavior per nest from the beginning, middle, and end of monitoring phase (mean = 2.82 days per nest, SD = 1.46, range: 1—16 days). For 33 out of 293 nest days, the thermocouple measuring the ambient temperature malfunctioned, so in these cases we imputed the ambient temperature from the known elevation of the nest and the significant relationship between ambient temperature and nest elevation (linear model, R^2^ = 0.89, p < 0.001). Prior to this, we imputed missing elevation from six out of 104 nests by consulting data on species elevational ranges across the Manu gradient that we had collected over eight years of netting, observations and nest searching (GAL, AM, SKR, unpublished data).

We calculated two complementary metrics of nest attentiveness which are commonly used in the literature: diurnal and whole-day nest attentiveness. Whole-day nest attentiveness was calculated as the sum of minutes during which the incubating bird was on the nest divided by the total number of minutes during the day (thereby including nocturnal incubation bouts). We calculated diurnal nest-attentiveness as the sum of all diurnal minutes during which the bird was incubating on the nest divided by the number of minutes of daylight between local sunrise and sunset. For each day of incubation behavior and field site, we calculated local sunrise and sunset using the suncalc R package (*72*), and then summed all daily incubation on-bouts and added them to the diurnal portions of incubation bouts that either began before sunrise and ended after sunrise and the diurnal portions of the incubation bouts that began before sunset and ended after sunset. Diurnal and whole-day attention metrics were highly positively correlated (Pearson’s correlation coefficient = 0.727, t = 17.947, df = 288, p < 0.0001). We therefore focus on the diurnal nest attentiveness in the main text, as there was generally little variation in local sunset and sunrise, as all field sites were close to the tropics (7.04° < latitude < -13.17°).

***1.4. Nest microclimate and cooling data.*** For each incubation recess, we also noted the temperature decrease in °C, along with its average ambient temperature during the incubation recess as well as its duration. We also noted the number of eggs present during each day of incubation, as well as their fresh mass. Egg mass was measured in the field with a high-precision digital balance (FlipScale F2, Phoenix, AZ, USA) with an accuracy of 0.05 g. The product of the mean fresh egg mass and the number of eggs per nest yielded the cumulative egg mass per nest. This yielded 5,746 incubation recesses from 256 days of incubation from 92 nests from 33 species that full data available.

***1.5. Comparative data on hummingbird life history and biogeography.*** We collected data on hummingbird incubation periods and egg mass from three separate sources: 1) a literature study encompassing 3,096 birds (*6*), 2) Birds of The World (*73*) and references therein, and 3) our own data. For all analyses, we focused on obtaining species-specific means and then assembled a unified database: we first started with the data table from 1), then added any species (rows) that had any incubation period or egg mass data present in 2) or 3) and then filled any missing data on minimum nest height from 3).

When processing data from 2), we followed methods from (*6*). Briefly, we collected incubation period data from the section “Reproduction”, and calculated, when necessary, the midpoint between the reported range of incubation period. We also recorded the minimum nest height when present. The following life-history data were taken from BirdLife International (<http://www.datazone.birdlife.org>): to differentiate between migratory and resident species, we lumped all movement strategies (short- and long-distance, altitudinal, and irruptive) into a single “migratory” category to differentiate these species from “resident” ones. We also noted the forest dependence of each species and scored it as (“high”, “medium”, “low” or “none”), following the methodology by (*6*). By visually inspecting the range maps from (*74*) we also categorized species into ones whose distributions were primarily insular or primarily continental, Body mass data was obtained from Avonet (*75*).

Using range polygons from BirdLife International (*74*) of the combined breeding and resident ranges, we calculated absolute latitudinal range midpoint, temperature and precipitation of the warmest quarter (bio10 and bio18) from WordClim (*76*) using the sf package in R (*77*, *78*).

***1.6. Comparative data avian egg surface area and volume.*** We extracted egg dimensions (average length, asymmetry and ellipticity) from a global analysis of 1,400 species (*41*). We first defined egg breadth (EB) as the fraction of average egg length (AEL) and egg ellipticity (EE). We then calculated egg volume (EV) as follows, using k = 0.51, following (*79*):

$$EV =k x AEL x {EB}^{2}$$

We then calculated egg surface area (ESA) as follows, assuming s = 4.835, following (*80*):

$$ESA=s^{2/3} x {EB}^{4/3}$$

… and obtained the egg surface-area-volume ratio as the fraction of ESA/EV. While we acknowledge that egg asymmetry is also important in obtaining precise estimates of egg surface area and volume, we used readily available metrics and interpret our derived estimates of EV and ESA as – at best – proxy variables for the true quantity of interest, which is the surface area-volume relationship. However, even in the most complex ESA/EV estimation procedures, which require more specified conicity indices and do not yield the phylogenetic coverage we are interested in for the present study, we note that the simple ratio of egg breadth to length has been demonstrated to have the strongest influence on ESA/EV (*81*). We therefore consider our proxy of ESA/EV as sufficient for the purposes of the present study.

***2. Hummingbird egg surface-area volume analyses***

Using the newly derived proxy variable for egg surface-area-volume ratio (ESA/EV, see section 1.6), we explored phylogenetic patterns of ESA/EV ratios across all Aves. Ten out of highest eleven species-level ESA/EV ratios belonged to hummingbirds. We also summarized the available data by avian families and calculated mean ESA/EV ratios and their standard errors. We then obtained a family-level egg-trait tree by pruning a summary tree from the Global Bird Tree with the Hackett backbone (*82*, *83*) to one species per sampled family from (*74*). Finally, we performed an ancestral state reconstruction of family-level mean ESA/EV using the *fastAnc()* function from the phytools package in R (*84*), for illustrative purposes (Fig 2A).

***3. Analyses of incubation behavior***

***3.1. Defining incubation fragmentation.*** For each day of measured incubation behavior, we counted the number of incubation recesses, as well as the mean duration of all incubation recesses and incubation on-bouts. These metrics were highly interdependent (Fig. 2B-D), so we reduced these to a single principle component, using unrotated principal component analysis (PCA) in the R package ‘psych’ (*85*). Before performing PCA all three variables were log-transformed, Box-Cox transformed and subsequently centered and scaled to reduce predictor correlation and satisfy the assumption of Normality. The three metrics had strong loadings on the first principle component (PC) axis, which captured over 93% of the total variance (Table S1). As low values in this PC were associated with few incubation recesses and long incubation on-bouts, and high values described incubation behavior with many frequent interruptions, we hereafter refer to this as “incubation fragmentation” (Fig. 2B-D).

***3.2. Main results.*** In order identify drivers of incubation behavior, we estimated Bayesian phylogenetic linear mixed models, hereafter BPLMM, in the R package MCMCglmm (*86*) to determine to what extent the average diurnal ambient temperature, the species’ body mass, and the survival estimate of each nest contributed to the incubation behavior that each bird displayed.

We fitted models with incubation fragmentation as the response variable, and ambient temperature, species body size, the interaction between ambient temperature and body size, and the daily survival rate (see section 7) as predictor variables (i.e. fixed effects). We also accounted for the repeated sampling of nests by adding a random-intercept term for nest and the temporal structure of sampling with an unstructured random effect variance covariance matrix. Moreover, we added a phylogenetically informed random-intercept for species-level variation (*87*). We used a variance-covariance matrix derived from a species-level summary tree of the global Bird Tree with the Hackett backbone (*82*, *83*) to inform random effect level covariance. All predictors were scaled and centered before fitting the model.

We fitted three independent chains for every BPLMM and assessed their convergence through Gelman-Rubin statistics < 1.1, effective sample sizes above 8,000 for every parameter, and visual inspection of trace plots. Each model chain was run for 100,000 iterations with a burn-in of 20,000, a thinning interval of 3,000, and uninformative priors that included inverse Wishart distributions with ψ = 1 and ν = 0.002 for the random effects. Variance inflation factors were also computed and determined to be < 1.5, which is well below the often-used cutoff of ten (*88*). The following is an example of the R code used to estimate these models:

inv.phylo <- inverseA(mytree, nodes="TIPS", scale=TRUE)

MCMCglmm(incubation_pc1~ avg.ext.temp * mass + dsr, family="gaussian",

random=~species+us(1+nest_day_rank_numeric):nest_ID,

ginverse=list(species=inv.phylo$Ainv),

prior=list(R=list(V=1,nu=0.002),

G=list(G1=list(V=1,nu=0.002),

G2=list(V=diag(2),nu=2))),

data=df,nitt=100e3,burnin=20e3,thin=3e1, verbose=T)

… where *mytree* is the variable containing our phylogenetic hypothesis, *incubation_pc1* is the variable containing daily incubation fragmentation estimates for each nest’s observed day of incubation behavior, *avg.ext.temp* is the variable containing the daily mean ambient temperature during incubation recesses, *mass* is the incubating species mean body mass and *dsr* is the daily nest’s survival rate estimated (section 7); *species* denotes the species to which each day of incubation behavior belongs, *nest_ID* denotes the nest ID, *nest_day_rank_numeric* is the index of nest days sampled within nests, and *df* is the name of our dataset. The results of the converged model are shown in Fig. 2E, Fig. 3H & Table S2.

***3.3. Results using nest attention.*** As comparative analyses of incubation behavior often use aggregate metrics of nest attentiveness (*12*, *25*) to describe variation in parental investment and adult behavior, we also calculated diurnal nest attentiveness as above (section 1.3). We show how diverse incubation strategies might fail to be distinguished by nest attentiveness (Fig. S1) and evaluated the correlation between nest attentiveness and incubation fragmentation in our dataset using Pearson’s correlation test (Fig. S2). We then repeated the analysis above (section 3.2.) using nest attentiveness as the response variable instead of *incubation_pc1* (Table S3).

***4. Robustness analyses of incubation behavior***

***4.1. Phylogeny in Principal Component Analysis.*** We used PCA to capture the three aspects of daily incubation behavior that we were interested in (incubation recess frequency, mean incubation recess duration, mean incubation on-bout duration). However, one assumption of PCA is that observations are independent of one-another, which is violated by comparative data, as shared ancestry makes some species-pairs more similar to each other than others (*84*). Although this assumption is violated by our dataset, it is additionally violated by repeated sampling of nests. To our best of knowledge, we are unaware of PCA routines that can simultaneously account for repeated sampling as well as shared ancestry. We therefore extended the phylogenetic PCA approach by (*84*) to account for repeated sampling of species. In this approach, we graft novel tips onto the phylogenetic species tree (which has one tip per species), such that the tree becomes a tree of samples, inspired by (*89*). We then pass this novel tree of samples to the pPCA routine, and hereafter refer to this technique as “hierarchical phylogenetic principal component analysis” (hpPCA).

The loadings from the hpPCA were similar to those of the standard PCA (Table S1) (Loadings on PC1 in hpPCA; log number of daily incubation recesses = 0.965, log mean incubation bout duration = -0.787, log mean incubation recess duration = -0.851). The first PC captured 73.5%, the second one captured 23.9% and the third captured the remaining 2.6% of the total variance. The first axis of the hpPCA was significantly positively correlated with the first axis of the PCA (Pearson’s correlation coefficient = 0.9999, t = 1218.5, df = 291, p-value < 0.000001). Rerunning the BPLMMs to analyze incubation behavior using the first axis from the hpPCA instead of the PCA yielded qualitatively identical conclusions (Table S4).

***4.2. Phylogenetic Uncertainty.*** We explored the influence of phylogenetic uncertainty by refitting the model of incubation behavior from the main analyses on 100 different phylogenetic hypotheses from the posterior distribution of the Global Bird Phylogeny from the Hackett backbone (*82*, *83*), allowing one chain to be run on each phylogenetic hypothesis and averaging all posterior results from the 100 chains/trees. The results were qualitatively identical (Table S5).

***5. Analyses of nest temperature cooling***

To explore how fast and why nests cool when left unattended by incubating adults, we used fine-scale data on incubation events, for which we had identified incubation recess duration, as well as average ambient temperature, and the drop in temperature over the course of the incubation recess. We also had data on each nest’s egg mass and clutch size, such that our full analytical data set consisted of 5,476 incubation recesses, from 256 days of nest observations, from 92 nests, covering 33 species (some nests lacked egg mass data). To this dataset we fitted multipredictor generalized additive models (GAMs) in which incubation recess duration had a nonlinear effect on the drop in nest temperature with three degrees of freedom.

***5.1. Nonphylogenetic analyses***

We first fitted nonphylogenetic, descriptive GAMs to our dataset, with a model structure as below, in which ‘*s()’* denotes a nonlinear effect, ‘*:’* denotes an interaction, and ‘*+’* denotes additive effects. We removed the intercept and all additive effects from this model to ensure that nest temperature cannot drop without an incubation recess of at least one minute. Herein, *temp_drop* describes the decrease in nest temperature over the course of the incubation recess, *Trip.Length* is the duration of the incubation recess, *avg.amb.temp* is the average ambient temperature during the incubation recess, and *egg_mass_cumulative* is the total fresh mass of eggs in that nest, and *off_bouts_clean* is our dataset. The following is an example of the R code used to estimate this model:

gam(temp_drop~0+s(Trip.Length)+ avg.amb.temp+

s(Trip.Length):avg.amb.temp+

s(Trip.Length):egg_clutch_mass_cumulative+

s(Trip.Length):egg_clutch_mass_cumulative:avg.amb.temp+

avg.amb.temp:egg_clutch_mass_cumulative,

data = off_bouts_clean)

Two degrees of freedom (k) were sufficient to describe the decrease in temperature that eventually approached ambient temperature as incubation recess duration increased, as we noted that model fit eventually plateaued after two degrees of freedom (Akaike Information Criterion (AIC) of interceptless model and nonlinear effect of incubation recess duration: AIC_k=1_ = 21,977.8, AIC_k=2_ = 21,540.8, AIC_k=3_ = 21,439.2). Given that the multipredictor model estimated the degrees of freedom for the non-linear effect to be equal to three, we report the model with three degrees of freedom. Moreover, simulating Newtonian cooling and subsequent GAMs with varying degrees of freedom revealed no substantive benefit of increasing degrees of freedom beyond three, and spurious increases in temperature at long incubation bout durations at two degrees of freedom (see code). We assumed that average ambient temperature modified the effect of extending incubation bout durations, and that increasing clutch mass could buffer temperature losses either by alleviating the effect of extended incubation recesses, alleviating the effect of cold temperatures, or alleviating the way cold temperatures exacerbate temperature loss during extended incubation recesses. The results of these descriptive GAMs are shown in Fig. Fig. 2J-K, Fig. S3 and Table S6. We chose to report this model in the main text as the interpretation of non-linear effects are straightforward and all portions of the nonlinear effect of incubation recess duration could be modelled as interactions.

***5.2. Phylogenetic analyses.*** We then fitted phylogenetic generalized additive mixed models, which incorporate phylogeny, repeated sampling, and non-linear effects using the MCMCglmm R package (*86*) to see how robust the descriptive model is. Herein, nonlinear effects must be fitted as separate components of a natural spline, which then requires a choice about which basis of the spline that additional effects should interact with: visual inspection of the spline basis loading revealed that the second spline basis loading on the variable describing incubation recess duration was highest (especially at short/intermediate incubation recesses, which were the most frequently observed incubation recesses: 95.3% of all were shorter than 20 minutes), which suggested that the modifying effects of ambient temperature and clutch mass should interact with the second spline basis function (see code). This choice was confirmed by the fact that the model had the strongest effect size for the second spline basis function for the duration of incubation recess (Table S7). Therefore, we fitted a model as below, with the same naming conventions as above; below is an example of the R code used to estimate this model.

MCMCglmm(temp_drop~0 +Trip.Length_spline1 +

Trip.Length_spline2 +

Trip.Length_spline3 +

Trip.Length_spline2:avg.amb.temp +

Trip.Length_spline2:egg_clutch_mass_cumulative +

Trip.Length_spline2:egg_clutch_mass_cumulative:avg.amb.temp +

avg.amb.temp:egg_clutch_mass_cumulative,

family="gaussian",

random=~species+nest.day,

ginverse=list(species=inv.phylo$Ainv),

prior=list(G=list(G1=list(V=1, nu=0.002),

G2=list(V=1, nu=0.002)),

R=list(V=1, nu=0.002)),

data=df,nitt=100e3,burnin=20e3,thin=3e1, verbose=T)

As each nest yielded multiple days of incubation behavior, we also accounted for repeated sampling of nests by adding a random-intercept term for “day of measurement within nest”. Moreover, we added a phylogenetically informed random-intercept for species-level variation. We used a variance-covariance matrix derived from a species-level summary tree of the Global Bird Tree with the Hackett backbone (*82*, *83*) to inform random effect level covariance. All predictors were scaled and centered before fitting the model.

We fitted three independent chains for every model and assessed their convergence through Gelman-Rubin statistics < 1.1, effective sample sizes above 8,000 for every parameter, and visual inspection of trace plots. Each model chain was run for 100,000 iterations with a burn-in of 20,000, a thinning interval of 3,000, and uninformative priors that included inverse Wishart distributions with ψ = 1 and ν = 0.002 for the random effects. As the model does not have an intercept, traditional VIFs were not used to evaluate collinearity. However, we suspect that collinearity was unlikely to be important, as pair-wise predictor correlations were low (ρ ≤ |0.433|). The results from the phylogenetic and non-phylogenetic generalized additive models were qualitatively similar (Fig. S4, Tables S6-7), although the support for the modulating effects of clutch mass and temperature were somewhat diminished. Nevertheless, this is to be expected to some degree, as only one portion of the non-linear effect of *Trip.Length* was allowed to interact with clutch mass and temperature as the modulating interactive effects are diluted across each of the three spline parameters, in contrast to the non-phylogenetic model which concentrates the modulation into single parameters (see above). As the effect sizes among the standard and phylogenetic models have the same polarity and have relatively similar magnitudes, we therefore conclude that both models support qualitatively similar descriptions of the system.

***6. Analyses of hummingbird life history***

To explore correlates of life history across all hummingbirds, we used phylogenetic generalized linear models to evaluate the factors associated with variation in incubation period and egg mass, with species-averages as the sampling unit. In all models, we accounted for shared ancestry by using a consensus tree derived from the Global Bird Tree with the Hackett backbone (*82*, *83*).

***6.1. Ecogeographic drivers of hummingbird incubation periods & egg mass.*** We first considered drivers of incubation period, which log-transformed prior to analysis. We used phylogenetic regression models in the R package *phylolm* (*90*) with Pagel’s lambda to assess the model of evolution. We employed backwards model selection to identify the final model, and as the variable of nest height had 35 missing values, we scrutinized that variable first and then simplified models by sequentially eliminating the variable with the largest p-value and refitting. We continued model simplification until only significant parameters remained. Variance inflation factors in the full model were below 4, well below the cutoff of 10 (*88*). All continuous predictors were Box-Cox-transformed, scaled and centered before model fitting. The results for incubation periods are shown in Table 1, Fig. 2L & Fig. 3J. We then repeated this procedure on the variable of (log-transformed) egg mass (Table S8).

***6.2. Relationship between incubation period and egg mass.*** In addition to evaluating the multiple traits of parental investment (incubation period and egg mass) separately, we also wanted to see if, as expected (*18*), longer incubation periods are facilitated by larger investment in egg mass. As both traits should have expected strong allometric relationships with body mass, we therefore considered only the three traits (body mass, incubation period and egg mass) in three additional models. First, we took a “two-step” approach, in which we fitted two separate log-log phylogenetic regressions of incubation period and egg mass against body mass, respectively. Then we extracted the residuals from these models (yielding relative egg mass and relative incubation period) and performed a subsequent phylogenetic regression of these two new metrics against one another.

As the use of residual traits has been criticized on statistical grounds (*91*, *92*), we therefore adopted a complementary second approach, in which we fitted two log-log phylogenetic regression models that also had body mass as a separate predictor (alongside egg mass or incubation period, respectively). We assessed the presence of correlated traits of parental investment based on 1) the effect of residual egg mass on residual incubation period, 2) the effect of egg mass on incubation period while accounting for body mass and 3) the effect of incubation period on egg mass while accounting for body mass (Fig. S5, Table S9).

***6.3. Robustness to phylogenetic uncertainty.*** We first explored the effect of phylogenetic uncertainty on the drivers of hummingbird life history. For models fitted to a single phylogenetic hypothesis in the *phylolm* R package (*90*), the functions available in the *sensiPhy* package (*93*) allow for the incorporation of multiple phylogenetic hypotheses. We expanded these functions to additionally allow for multiple predictors in phylogenetic regression models (following (*94*)) and refitted our final models of incubation period and egg mass on 100 additional phylogenetic hypotheses. The results remained qualitatively identical, although the support for the marginally significant tradeoff between residual incubation length and egg mass decreased (Table S10).

***7. Survival analyses***

We discovered 390 nests of the 34 target species for which we obtained detailed incubation behavior. On average, each species was represented by 11.4 nests (SD = 11.8, range: 1 – 49). Of these nests, 117 were successful, 178 were depredated, 84 failed and 11 had unknown nest fates (i.e. the survey period ended before nest fate could be ascertained). This generated 6,092 days of nest-observation days. On average, each nest yielded 15.6 (SD=11.5, range: 2-53) days of observation. When nests were discovered with nestlings, we conservatively assumed that the nest was also active at least 1 day before the nest was found. For each nest, we calculated the total number of observation days as the number of days that elapsed from the day of nest discovery until the final day of nest observation. Finally, we also pooled all observation days and nest fates from all nests to estimate daily nest survival and predation rates using the Mayfield estimator (*45*), pooling all nests first across and within field sites (Fig. 3AB, Table S11).

***7.1. Modelling daily nest survival rate.*** Both analyses of daily nest survival rate and daily nest predation rate (section 7.2.) were performed analogously, using phylogenetic logistic exposure models (*46*), using the number of observation days as an offset and a binary response variable. For nest survival, we used nest fate (1=success/unknown, 0=predation/failure) as a response variable to estimate nest survival as a function of the predictor variables. For nest predation, we used nest predation (1=predation, 0=no predation) as a response variable to estimate nest predation rate as a function of the predictor variables, which were: nest height (cm above ground), elevation (m above sea level), interannual variation (year as a categorical effect), and species-specific variation. Specifically, we allowed for phylogenetically pooled random intercepts ((*87*)), in which species-level deviations from a shared intercept were informed by shared ancestry, assuming a species-level consensus tree from BirdTree assuming the Hackett backbone (*82*, *83*), which we constructed following (*94*). As intraspecific variation in nest height was substantial and well-sampled (mean intraspecific SD in nest height =78.6), we also allowed for species-specific effects of nest-height (i.e. phylogenetic random slopes of nest height). As only few species were found across multiple stations due to strong elevational replacement in hummingbirds (*95*, *96*), we fitted a community level effect of elevation as in (*42*).

We fitted the phylogenetic logistic exposure models in brms (*97*–*99*), running the models with three chains, for 4,000 iterations with 2,000 samples as warm-up each. We set adapt_delta=0.95, and scaled and centered predictors to improve chain mixing. We used weakly informative priors (Normal distributions with mean zero and standard deviation 2) on fixed effects and vague priors (Student’s T-distributions with mean zero, 3 degrees of freedom and scale = 2.5) on random effects. All model parameters converged with Gelman-Rubin statistics < 1.1 and ESS > 1,000 and passed visual inspection of trace plots. All code is provided in Supplementary Data. The following is an example of R code used to estimate this model.

brms_model_phylo <- brm(

nest_fate ~ elevation + height +year + offset(log(days_observed)) +

(1+height | gr(species, cov = vcv_matrix)),

data = df,

family = bernoulli(link = "logit"),

data2 = list(vcv_matrix = inv.phylo),

prior = c(

prior(normal(0, 2), class = "b"),

prior(normal(0, 2), class = "Intercept"),

prior(student_t(3, 0, 2.5), class = "sd")

),

chains = 3, cores = 3, iter = 4000, warmup = 2000, control = list(adapt_delta = 0.95)

)

…in which *nest_fate* denotes binary nest survival, *elevation* denotes the elevation of the nest, *height* denotes the height of the nest above the ground, *days_observed* the number of days that each nest was observed for, *df* denotes the data table and vcv_matrix shows the variance scaled covariance matrix derived from the species-level phylogeny. The results are shown in Figure 3C-H and Table S12.

***7.2. Drivers and consequences of daily nest predation rate.*** Our phylogenetic logistic exposure models utilized all sources of nest mortality and thus estimated daily survival rate. We therefore reran the logistic exposure models of daily survival rate above using only the 178 nest failures due to predation and inverting the Bernoulli response variable (1=nest predation, 0=no nest predation). We retain identical predictor variables and run conditions, and obtained qualitatively identical models of drivers of daily predation rates (Fig. S6, Table S13)

To explore if daily predation rate explained variation in incubation behavior, we reran the models of incubation behavior (section 3.2), substituting daily predation rate for daily survival rates. The results were qualitatively identical (Table S14).

***7.3. The effect of survival on incubation period.*** To explore the role of survival on incubation periods, we consulted an additional database of avian survival rates worldwide (*39*). Of the 147 species we had data on incubation period, only 33 species also had survival rates available. Although the database of survival rates offers multiple samples per species, for the species of interest in our study, only limited repeated sampling was present (one species had three samples, three species had two samples, and the remaining 29 species had one sample each). Therefore, we averaged the survival estimates by species for the four species with multiple samples and used phylogenetic regression models on species means as above. We performed backwards model selection, including survival rate as a covariate in the full model. However, this model had substantially lower sample size (N = 22) compared to the full model of the previous analysis. We then performed model selection, first evaluating the two variables with the most restrictive sample sizes (survival rate and nest height), and then sequentially removing variables according to their p-values (Table S15). To be completely sure that survival rate had no effect on incubation period, we refitted the final model (which had only body mass, temperature and insularity) by adding survival rate, but this had no significant effect, thereby repeatedly justifying its elimination (estimate = 0.010, SE = 0.026, t-value = 0.387, p-value = 0.703, Fig. 3I).

***7.4. Robustness to measurement error.*** As our comparative estimates of daily survival rate also included standard errors (*34*), we then explored the effect of measurement error on the effect of daily survival rate on hummingbird incubation period. Specifically, we fitted a Bayesian phylogenetic metaregression model with a measurement error (*100*) on the term of daily survival rate that included the standard errors included in the original data source to explore possible drivers of variation in hummingbird incubation period. The model code is shown below:

brm(bf(log_incubation_d ~ me(DSR, DSR_se) + log_body_mass +

temperature + log_precipitation + log_latitude +

log_nest_height_min + habitat + migration + insularity +

(1|gr(phylo, cov = A)) + (1|species)),

data = inc_dur_analytic, family = gaussian,

iter = 600000, warmup = 100000, thin=500, cores = 3, chains = 3,

data2 = list(A=A), control = list(adapt_delta=0.9))

… where “*DSR*” and “*DSR_se*” indicate variables containing the means and standard errors for the daily survival rates from (*39*), and “*inc_dur_analytic*” is the name of our dataset and *me()* indicates that a measurement error model is fitted for that predictor. Our results confirmed that daily survival rate continues to have no effect on incubation period in hummingbirds, thereby justifying its removal in model selection (Table S16).

We fitted the phylogenetic metaregression model in brms (*97*–*99*), running the model with three chains for 600,000 iterations and 100,000 samples as warm-up and thin rate of 500. We set adapt_delta=0.90, and scaled and centered predictors to improve chain mixing. We used weakly informative priors (Normal distributions with mean zero and standard with standard priors. All model parameters converged with Gelman-Rubin statistics < 1.1 and ESS > 1,000 and passed visual inspection of trace plots.

***8. Validation of Assumptions***

***8.1. Relationship between ambient temperature and incubation fragmentation.*** We assumed that temperature and incubation fragmentation are related in all hummingbird species outside of our sample. While we did not have detailed incubation behavior data on all 147 species of hummingbirds, we were able to evaluate this assumption of the relationship between incubation fragmentation and incubation recess number in a sample of 201 species of birds worldwide (*36*), from which we obtained species mean estimates of daily incubation recess durations and recess frequencies. We then defined a univariate aggregate metric of incubation fragmentation as above, using principal component analyses in psych, after log-transforming, scaling and centering the variables. This aggregate metric (PC1), captured 94% of total variance, and had similar loadings as the incubation fragmentation metric as above (PC1 loading offbout duration = -0.97; PC1 loading offbout frequency = 0.97). Using phylogenetic PCA (pPCA, (*84*) instead of standard PCA yielded similar loadings (pPC1 loading offbout duration = -0.94; pPC1 loading offbout frequency = 0.78, pPC1 captures 77.0% of total variance) and near identical values of PC1 (PC1 ~ pPC1, correlation coefficient = 0.9989, p < 0.0001), such that we refer interested readers to the code supplement.

We then quantified the species-wide mean ambient temperature during the breeding season using range maps (*74*) and the mean temperature during the warmest quarter (“bio10” WorldClim (*76*)), as above (section 1.5). Next, we used a species-level tree from with the Hackett backbone (*82*, *83*) to perform phylogenetic regression of incubation fragmentation against mean ambient temperature using phylolm and Pagel’s model of evolution (*90*) (Fig. S8).

***8.3. Exploring Possible Target Temperatures.*** In Figure 4C-F, we present example temperature traces of one single day each of nests of species with low and high incubation fragmentation. To illustrate how cumulative daily exposure to target incubation temperatures could vary across incubation strategy, we selected all nest temperature measurements that were made during diurnal incubation bouts and fitted Gompertz models in the R package ‘thermPerf’ to describe the nonlinear warming (*101*). Gompertz functions are widely used to model biological growth (*102*) and return a biologically relevant asymptote parameter. For each nest, we fitted curves and obtained the asymptote parameter, and multiplied this by 0.95, to obtain a plausible nest temperature that could correspond to the target incubation temperature, based on the observation that birds stop increasing nest temperature when this value is surpassed. While these values differ between these two nests, we do not make any assessment of our unmeasured egg temperatures. To visually illustrate how incubation behavior influences the difference between realized and target temperatures, we selected three days of incubation across the elevational gradient, and chose plausible nest temperatures that correspond to target temperatures for graphical display (Fig. S9).

***8.3. Relationship between nest and egg temperature.*** We assumed that nest temperatures, which we measured in all 104 nests, approximated egg temperatures. In hummingbirds, measurement of egg temperatures is technically challenging, as the eggs are so small that inserting a thermocouple is exceedingly difficult. We do not attempt to estimate egg temperature from nest temperature, as small changes in positioning of nest sensor location can change absolute values of nest temperature, and a multitude of factors such as ambient temperature, nest insulation, nest microclimate, adult behavior and posture, adult body temperature, clutch size, etc. could all modify any structural relationship between nest and egg temperature. Nevertheless, we were able to validate our assumption that nest temperature was significantly positively related to egg temperature based on a limited sample of seven total days of incubation from two nests (one Green Hermit, *Phaethornis guy*, and one unidentified hermit, *Phaethornis sp.*). To measure egg temperatures, we sacrificed one egg and inserted a thermocouple, sealing the hole with superglue and attaching the temperature sensor to the same dataloggers as above. Intra-day correlations between nest and egg temperature were highly significant and positive (range: 0.60 < ρ < 0.985; Figure S11).

***8.4. Nocturnal Body Temperatures.*** Target egg temperatures during incubation are constrained by adult body temperature. Barring direct measurements of hummingbird egg temperatures across the elevational gradients studied here, we turned to a previously collected dataset on hummingbird internal body temperatures (*58*) that were collected at three field sites in Colombia between 520 and 2,500m elevation for a study of naturally occurring torpor (see paper for detailed methods). Briefly, internal body temperatures of captive adults at ambient temperature were measured with cloacal thermistors attached to U-12 HOBO data loggers (Onset Computer Corporation, Pocasset, MA, USA, <https://www.onsetcomp.com>), programmed to take 1 temperature measurement per minute. Each data logger was connected to 2 thermocouples, one of which was inserted into the experimental birds’ cloaca, the other measured ambient temperature. As torpor usage was unambiguous but variable (48% of birds used torpor), we selected only the non-torpid birds’ temperature traces to estimate mean nocturnal body temperature to see if this varied systematically across the elevational gradient. Moreover, to avoid any body temperature variation due to handling stress, we selected only the portion of each birds’ body temperature that was measured between 8pm and 4am local time. This yielded 112 estimates of mean adult nocturnal body temperature from 24 species.

We then explored if mean nocturnal body temperature varied with elevation by fitting a phylogenetic generalized linear mixed model in brms (*97*–*99*), using a phylogenetic tree from the Global Bird Phylogeny using the Hackett backbone (*82*, *83*). The model code is shown as follows:

brm(data = df, family = gaussian,

bf(mean_body_temp~elevation_scale+(1|gr(phylo, cov = A)) + (1|species)),

iter = iter, warmup = warmup, thin=thin, cores = 3, chains = 3,

data2 = list(A=A),

control = list(adapt_delta=0.95))

… where “*mean_body_temp*” indicates the mean nocturnal internal body temperature, “*elevation_scale”* is the centered and scaled variable of station elevation where birds were caught and “*df*” is the name of our dataset.

We ran the phylogenetic generalized linear mixed effects regression model with three chains for 600,000 iterations and 200,000 samples as warm-up and thin rate of 100. We set adapt_delta=0.95 to improve chain mixing. We used weakly informative priors (Normal distributions with mean zero and standard with standard priors. All model parameters converged with Gelman-Rubin statistics < 1.1 and ESS > 1,000 and passed visual inspection of trace plots (Table S17), and elevation had no significant effect on nocturnal body temperature (Table S17, Fig. S12).

Full Acknowledgements

We thank Dr. Jill Jankowski and everyone on the Manu Bird Project. The following field technicians helped find the 390 hummingbird nests that were used in this paper: Audrey Adams, Richard Aracil, Nestor Arilio Peralta, Nik Aspey, Clifton Avery, JWB, Katie Becraft, Sinead Borchert, Cornelio Bota-Sierra, James Brennan, Alejandro Campuzano, Adam Carter, Juliana Ceron Cardona, Julio Cesar Bermudez, Andres Chinome, Miranda Ciotti, Santiago David Rivera, Justin Demaniew, Juan Diego, Jacob Drucker, Matthew DeSaix, Jonny Alexander Echeverri Calderón, Maria Camila Estrada Florez**,** Colin Fagan, Wieland Feuerabendt, Eric Fischel, Samuel Flake, Camilo Florez, Michael Fuss, Dano Grayson, Diego Guevara Torres, Dave Gio, Jaime Garizabal, Naman Goyal, Rachel Hanauer, Jenny Hazlehurst, Erin Johnston, Michele Kelly, Jeremiah Kennedy, Callum Kingwell, Catherine Klein, MALM, Gliselle Marin, Anthony Miller**,** Jenny Muñoz, Simon Nockold, Sebastian Perez Peña, David Ocampo Rincon, Diego Rinconguarin, JSH, Jhan Carlos Salazar, Manuel Sanchez Martinez, Corey Shake, Giovanny Valencia, Wendy Valencia, Wendy Vidal Hernandez, Jesse Vooz & Cynthia Zurita Cervera. Lina Peña Ramirez provided photos and assisted with figure design in Figs. 1, 2 and S2. Isabella Burgos designed the icon in Fig. 1A. We thank the following photographers for sharing their photos on Wikimedia Commons with Creative Commons licenses: Charles J. Sharp, Félix Uribe, Mike Baird, Hector Bottai, Melissa McMasters, Wikidieren and Christoph Moning. We thank Ferran Sayol, Margot Michaud and Anthony Caravaggi for contributing icons to PhyloPic. We thank Carlos Calle for permission to use the photo of *Phlogophilus harterti*. We thank Emily Wheeler for editorial assistance, Lina Peña Ramirez, Jorge Lizarazo and I. Baldwin for discussions. We thank BirdLife International for access to range maps. The assistance of Marianne van Vlaardingen (Pantiacolla Lodge), Daniel Blanco (Cock-of-the-Rock Lodge), the Wayqecha Cloud Forest Biological Station, Gustavo Campuzano (Remedios), Michelle Tapasco (Montezuma Rainforest Lodge, Tatamá), La Minga Ecolodge and Icesi University (Zygia Biological Station) was invaluable. We thank SERNAP for permission to work in the Manu Park buffer zone in Peru and PNN Farallones de Cali and PNN Tatamá in Colombia for permits.

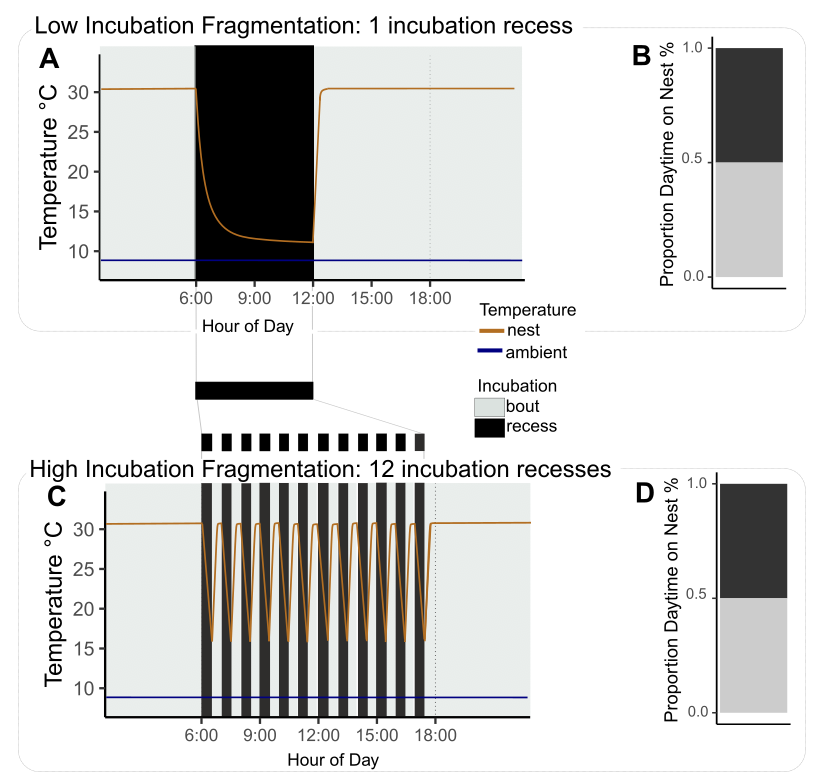

**Fig. S1. Nest attentiveness fails to capture important variation in incubation behavior.** Cartoon example of two incubation strategies with either high (**A-B**) or low incubation behavior fragmentation (**C-D**). (A) Incubation recesses are condensed into a single six-hour trip away from the nest and adults spend one entire six-hour period during daylight hours on the nest. (B) This yields 50% diurnal nest attention (6h each on/off nest during daylight hours). In contrast, when species fragment their diurnal incubation behavior into 24 incubation bouts and recesses of 30 minutes each (C), they can still reach 50% diurnal nest attentiveness (D). Nest temperatures (orange lines) were simulated using exponential decay and logistic growth functions. Code is provided in supplement. In A and C, horizontal dark blue line denotes constant ambient temperature.

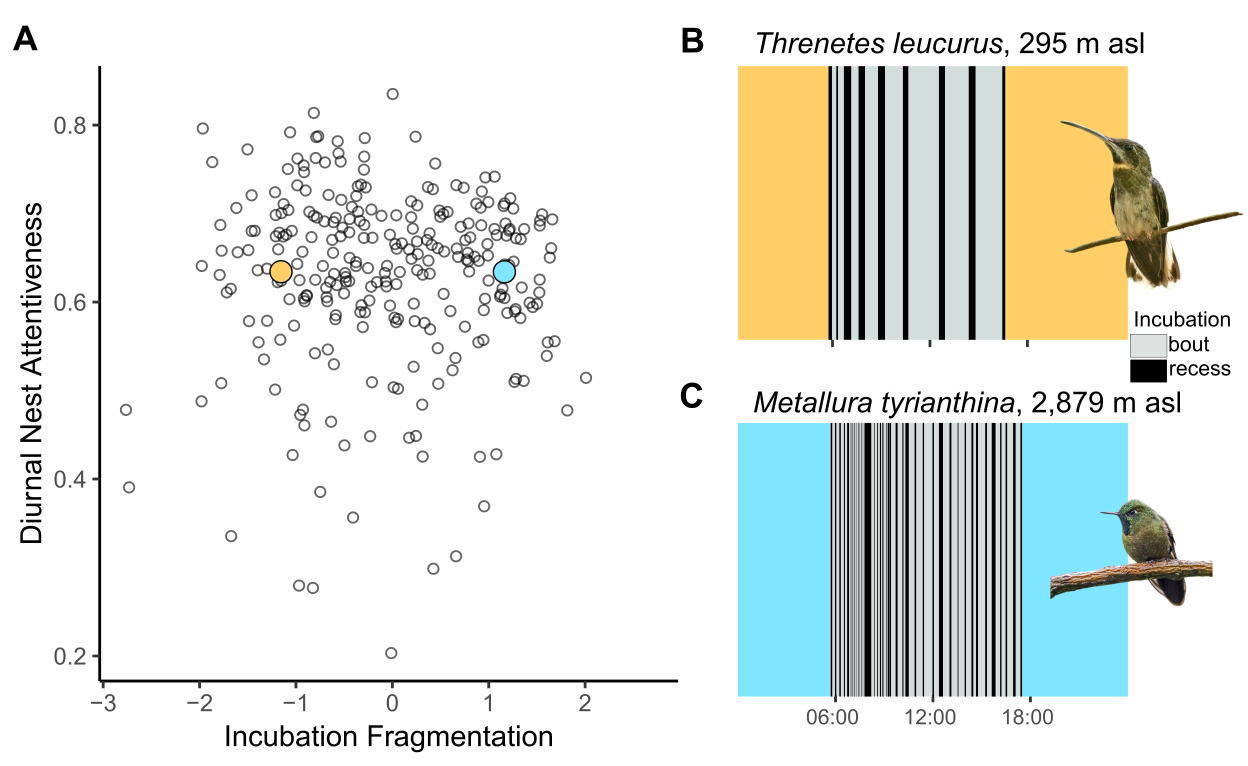

**Fig. S2. Diurnal nest attentiveness is unrelated to incubation fragmentation.** Incubation fragmentation reflects the daily number of incubation recesses, their duration, and the average incubation on-bout duration. Diurnal nest attentiveness is the proportion of minutes between sunrise and sunset that the parent bird spent on the nest incubating the egg. (**A**) The two metrics are not correlated (Pearson’s correlation coefficient = -0.015, t = -0.254, df = 275, p-value = 0.799). Blue point in (A) and (**B**) shows a day of incubation from a nest of *Metallura tyrianthina* at 2,879 m asl, red point in (A) and (**C**) shows a day of incubation from a nest of *Threnetes leucurus* at 295 m asl. Both days have diurnal nest attentiveness values of 63.4%, but incubation fragmentation ranges from -1.155 (*Threnetes*, B) to 1.162 (*Metallura*, C). Grey and block rectangles indicate incubation on-bouts and recesses, respectively. Photos provided by Charles J. Sharp and Hector Bottai from Wikimedia Commons and reproduced under CCSA4 licenses.

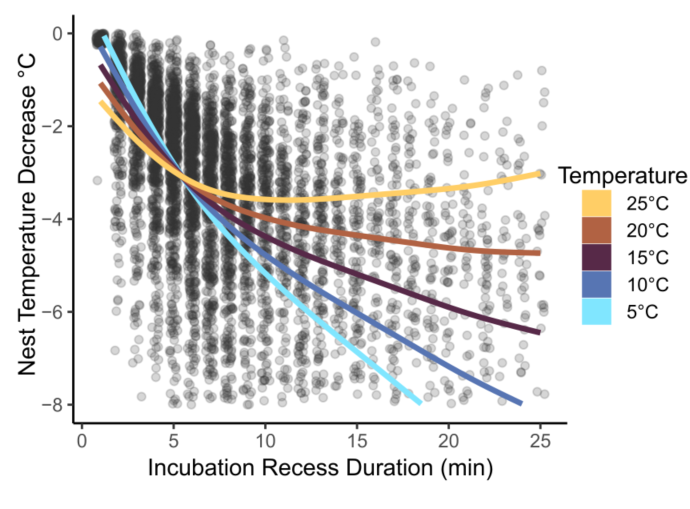

**Fig. S3. Nest temperature drops during incubation recesses and is modulated by ambient temperature.** Incubation recess duration has a non-linear effect on nest temperature decrease (natural spline base 3) in generalized additive of 5,476 incubation recesses over 256 days from 92 nests of 33 species of hummingbirds. Recess duration had strong effects in cold temperatures (interaction between incubation recess duration and ambient temperature: F_3513,1_ = 1480.018, p-value < 0.0001).

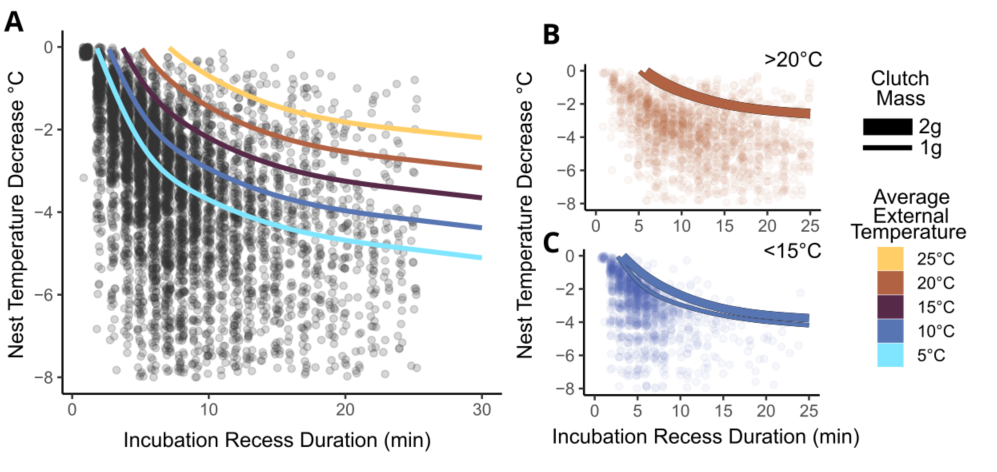

**Fig. S4. Nest temperature drops during incubation recesses and is modulated by clutch mass and ambient temperature in phylogenetic nonlinear mixed regression models.** Incubation recess duration has a non-linear effect on nest temperature decrease (natural spline base 3) in phylogenetic generalized additive mixed models of 5,476 incubation recesses over 256 days from 92 nests of 33 species of hummingbirds. Recess duration interacted with ambient temperature (**A**). Incubation recesses during warm conditions had similar effects for large and small clutch masses (**B**). When temperatures were cold (10°C), temperature loss was steep for small clutch masses and ameliorated for large clutch masses (**C**). Specifically, a 4°C drop in nest temperature in cold temperatures occurred in 20.7 minutes for 1g clutches but took 30 minutes to drop that low for 2g clutches.

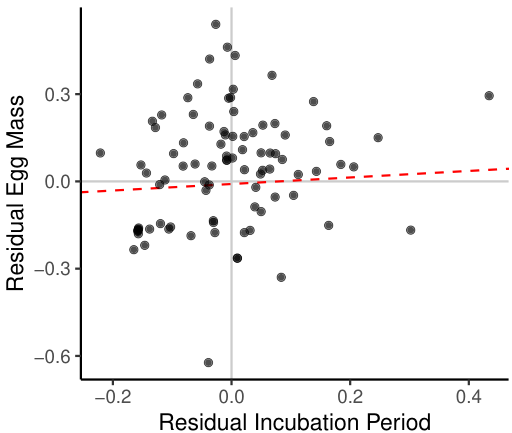

**Fig. S5. Weak positive relationship between egg mass and incubation period.** Phylogenetic regression model reveals marginally significant, positive relationship between residual egg mass and residual incubation period (estimate = 0.112, SE = 0.060, t = 1.878, p = 0.064, N = 89, R^2^ = 0.028, λ < 0.001). Residual egg mass and incubation period were estimated from two separate log-log phylogenetic regressions of egg mass and incubation period against body mass, respectively (egg mass-body mass allometry: estimate = 0.819, SE = 0.059, t = 13.971, p < 0.001, N = 89, R^2^ = 0.688, λ = 0.607; incubation period-body mass allometry: estimate = 0.336, SE = 0.023, t = 14.530, p < 0.001, N = 89, R^2^ = 0.671, λ = 0.562).

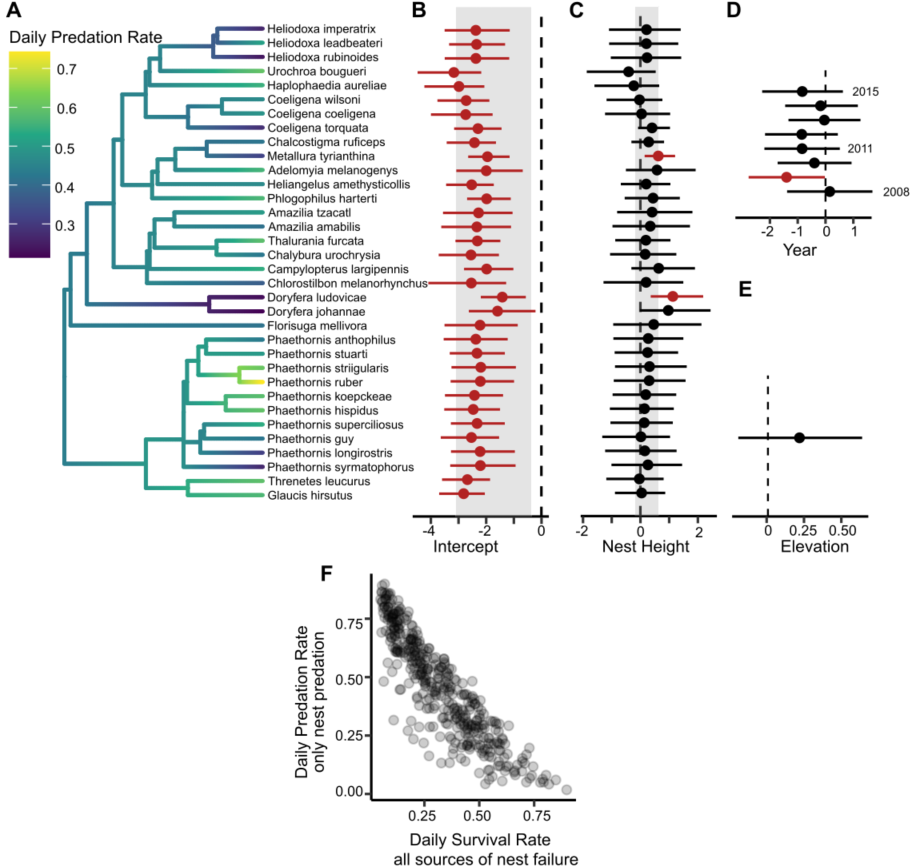

**Fig. S6. Nest predation rates of montane hummingbirds vary with phylogeny, not ecology.** Phylogenetic logistic exposure regression models reveal notable variation in daily predation rates across species, with weak species-specific effects of nest height. Vertical dashed lines indicate zero. (**A**) Tip color denotes species-average survival estimate of predicted values for each nest. (**B-E**) Points show species-specific posterior means and horizontal lines show 95% credible intervals of effect sizes from phylogenetic logistic exposure models, with significant effect sizes in red. Shaded rectangles show hyperparameter distributions for random intercepts (B) and random slopes of nest height (C). Effects of interannual variation (D) and elevation (E) applied to all species equally. (**F**) Daily survival rates (Fig. 2) are negatively correlated with daily predation rates.

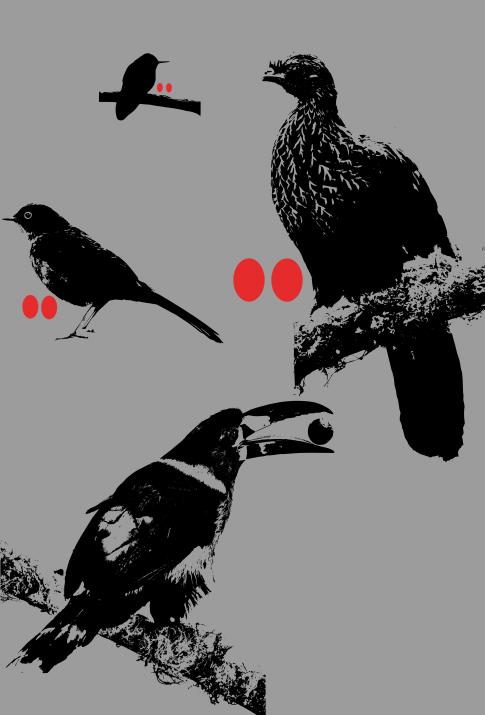

**Fig. S7. To-scale silhouettes of hummingbird, thrush, and guan and their eggs compared to a common high-elevation predator (Grey-breasted Mountain Toucan – *Andigena hypoglauca*).** Nest predators might gain less nutritional benefit from depredating hummingbird nests and consuming eggs (red) compared to e.g. thrush or guan eggs due to their tiny size. At high elevations in the Peruvian field site (Wayqecha), hummingbird nests (e.g. *Metallura tyrianthina*) contain two tiny eggs (mean mass = 0.504 g, SE = 0.014, N = 49), whereas thrush nests (e.g. *Turdus fuscater*) contain two medium eggs (mean = 10.5 g, SE = 0.114, N = 86) and guan nests (e.g. *Peneleope montagnii*) contain two large eggs (mean = 73.4 g, N = 1; all egg mass unpublished data from Wayqecha 2007-2014). Assuming equal nutritional content, depredating one *Turdus fuscater* nest delivers the same prey mass as depredating 20.8 *Metallura tyrianthina* nests, and depredating one *Peneleope montagnii* nest delivers the same prey mass as depredating 145.6 *Metallura tyrianthina* nests. Photos provided by Charles J. Sharp, Felix Uríbe and Wikidieren from Wikimedia Commons and reproduced under Creative Commons licenses.

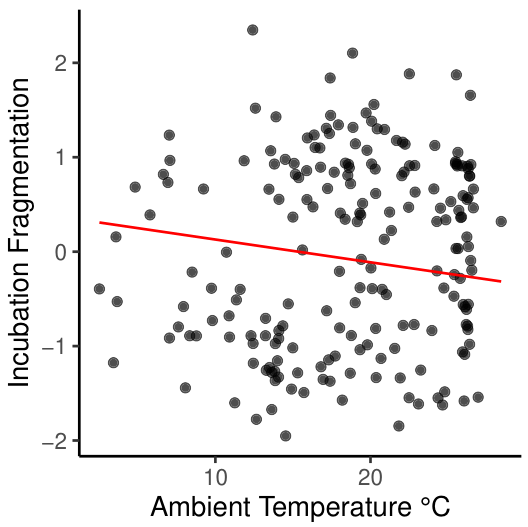

**Fig. S8. Temperature drives incubation behavior in 201 species of birds worldwide.** Phylogenetic regression of incubation fragmentation (PCA axis 1 of bivariate space of log incubation recess duration and frequency) on range-wide mean ambient temperature of warmest quarter (WorldClim variable “bio10”) reveals similar effect in 201 species worldwide (*36*) as in hummingbirds. Red line shows predicted relationship from phylogenetic regression (slope estimate = -0.146, SE = 0.037, t-value = -3.950, p = 0.00001, λ = 0.989, N = 201).

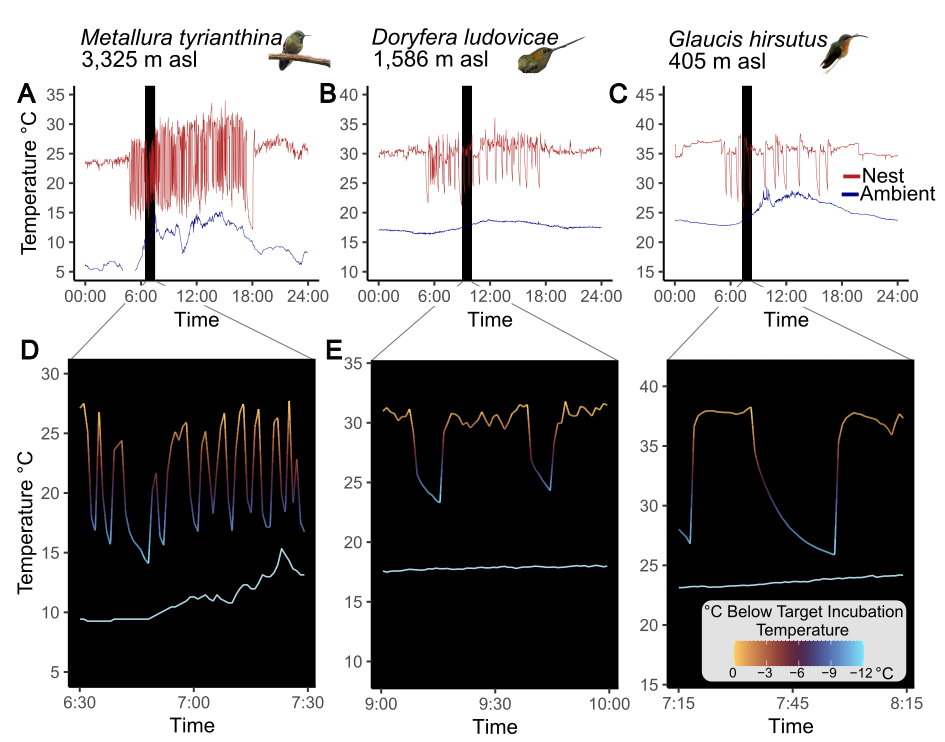

**Fig. S9. Nest temperatures warm asymptotically and cool rapidly, such that embryonic development efficiency could decrease as soon as adults leave the nest.** Temperature traces from whole days of incubation of three nests (**A**, *Metallura tyrianthina* at Wayqecha Biological Station in Peru at 3,325 m asl, **B**, *Doryfera ludovicae* at San Pedro Lodge in Peru at 1,586 m asl**, C** *Glaucis hirsutus* at Pantiacolla Lodge in Peru at 405 m asl). Dark blue lines show ambient temperature and dark red lines show nest temperature. Nest temperatures from one hour each (**D-F**). The target temperature during incubation could correspond to a specific nest temperature (visually estimated for each panel of where asymptotic warming could occur during each incubation on-bout, 27, 31, 38 °C respectively). Light blue line shows ambient temperature. Photos provided by Charles J. Sharp on Wikimedia and Lina Peña Ramirez.

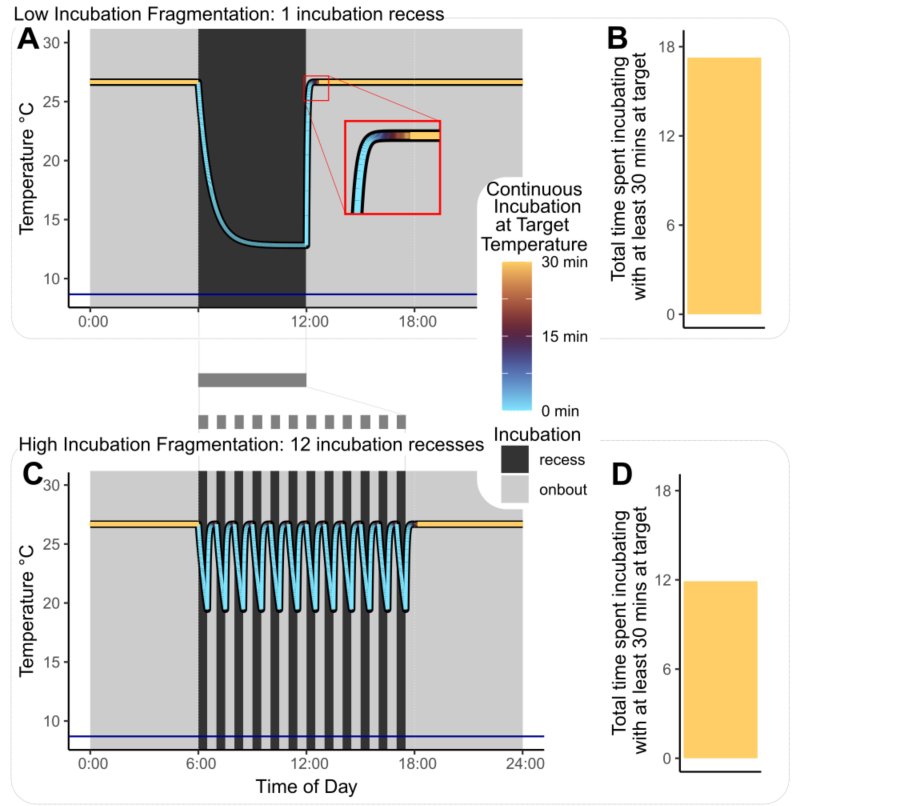

**Fig. S10. Incubation Fragmentation captures cumulative exposure to incubation at target temperature. (AB)** Cartoon example of two incubation strategies with either low (**A-B**) or high incubation fragmentation (**C- D**). (A) Incubation recesses are condensed into a single six-hour trip away from the nest and adults spend one entire six-hour period during daylight hours on the nest. (B) This yields almost 18 hours of daily incubation of at least 30 minutes continuous incubation at the target temperature at which embryonic development is most efficient. In contrast, when species fragment their diurnal incubation behavior into 24 on-bouts and recesses (C), they only achieve ca. 12 hours of daily incubation of at least 30 uninterrupted minutes at the target temperature (D), as diurnal incubation bouts are interrupted as soon as they reach 30 minutes duration. Nest temperatures were simulated using exponential decay and logistic growth functions. Code is provided in supplement. In A and C, horizontal dark blue line denotes constant ambient temperature.

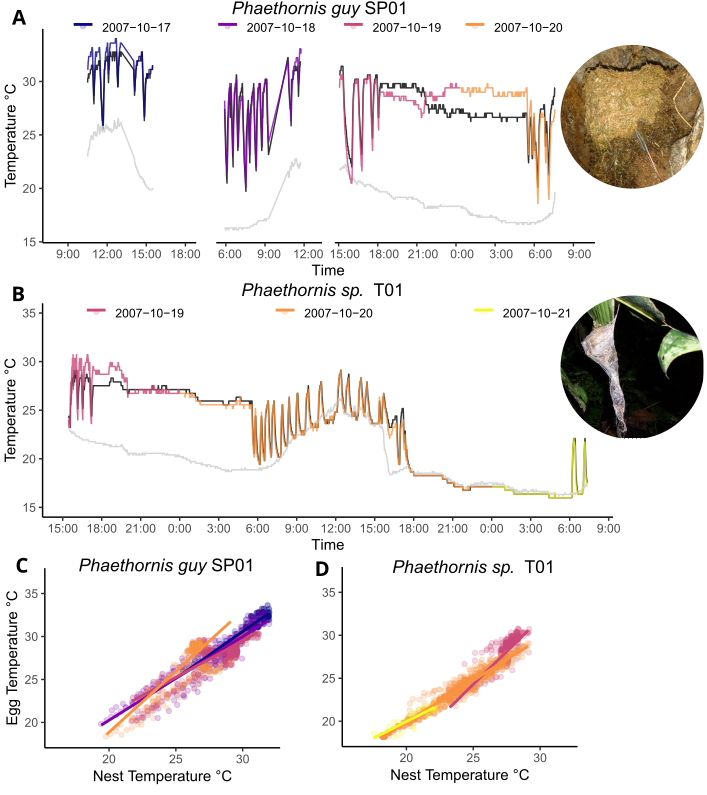

**Fig. S11. Egg temperature is significantly related to nest temperature in two hermit nests (*Phaethornis sp.*). (AB)** Egg and nest temperatures are highly related, with data from two nests which had thermocouples inserted into eggs (colored lines) and nests (black lines) and ambient temperatures (grey). Data were obtained from five unique days from two nests, one at San Pedro Lodge in Peru (A,C) and one at Tono River in Peru (B,D). Within-nest correlations in each nest (**CD**) range from 0.600 (2007-10-19 in SP01) to 0.985 (2007-10-20 in T01) and were all highly significant (P < 0.001). Photos by GAL. Mean diurnal ambient temperatures were 18.7° C at San Pedro Lodge and 20.2° C Tono River.

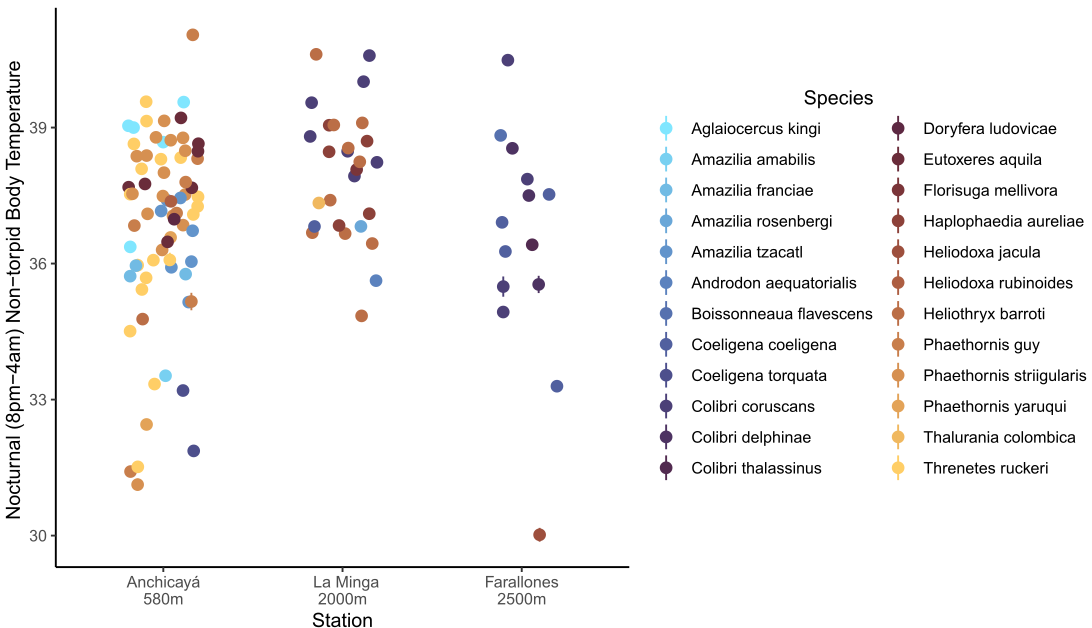

**Fig. S12. Adult internal nocturnal body temperature does not vary across elevation.** Adult mean body temperature during the hours of 8pm and 4am local time was calculated for 112 non-torpid individuals from 24 species of hummingbirds from three field stations in the western Andes (*58*). Points show means, and vertical bars show 95% confidence intervals. A phylogenetic mixed regression model revealed no significant effect of elevation on mean body temperature (elevation posterior mean = 0.21, SE = 0.52, 95% credible interval = 0.10-2.08).

**Table S1.** **Loadings of unrotated principal component analysis of daily incubation behavior.** Analysis conditions on a single axis of principal components and captures 93% of total variance of natural-log transformed, scaled and centered data.

| Variable | Incubation Fragmentation (PC1) |
| --- | --- |
| log Number of daily incubation recesses | 1.00 |
| log mean Incubation recess duration | -0.94 |
| log mean Incubation bout duration | -0.96 |

**Table S2. Temperature and body mass, not survival, influence hummingbird incubation behavior.** Phylogenetic generalized linear mixed models of incubation fragmentation. Daily survival rates were estimated for each nest in the prior logistic exposure models. Models converged with 3 chains, 100,000 samples, 20,000 burn-in, and thinning rate of 30. Within the G-structure, phylogeny has a posterior mean of 0.378, with effective sample size = 8,001. The R-structure (variable effects) has a posterior mean of 0.087, with effective sample size = 8,001. Significant parameters in bold, x denotes interaction.

| Parameter | Posterior Mean | Standard Error | 95% Credible Interval | | | Effective Sample Size | pMCMC |
| --- | --- | --- | --- | --- | --- | --- | --- |
| *Fixed Effects* |  |  |  |  |  |  |  |
| **Ambient Temperature** | -0.486 | 0.088 | -0.657 | — | -0.314 | 8,001 | **<0.0001** |
| Body Mass | 0.0003 | 0.104 | -0.205 | — | 0.199 | 8,652 | 0.9989 |
| **Daily Survival Rate** | 0.144 | 0.062 | 0.021 | — | 0.263 | 8,380 | **0.0217** |
| **Ambient Temperature x Body Mass** | -0.166 | 0.081 | -0.322 | — | -0.006 | 8,001 | **0.0432** |
| *Random Effects* on Nest ID | |  |  |  |  |  |  |
| σ Intercept | 0.334 | 0.086 | 0.193 | — | 0.528 | 8,289 |  |
| σ Day | 0.069 | 0.014 | 0.046 | — | 0.097 | 8,385 |  |
| ρ (Intercept, Day) | -0.078 | 0.030 | -0.145 | — | -0.028 | 8,305 |  |

**Table S3. Using diurnal nest attentiveness fails to reveal a dominant effect of temperature in driving incubation behavior.** Phylogenetic generalized linear mixed models of diurnal nest attentiveness. Daily survival rates were estimated for each nest in the prior logistic exposure models. Models converged with 3 chains, 100,000 samples, 20,000 burn-in, and thinning rate of 30. Within the G-structure, phylogeny has a posterior mean of 0.002, with effective sample size = 5,420. The R-structure (variable effects) has a posterior mean of 0.003, with effective sample size = 8,001. Interaction denoted by ‘x’.

| Parameter | Posterior Mean | Standard Error | 95% Credible Interval | | | Effective Sample Size | pMCMC |
| --- | --- | --- | --- | --- | --- | --- | --- |
| *Fixed Effects* |  |  |  |  |  |  |  |
| Ambient Temperature | 0.008 | 0.018 | -0.029 | — | 0.043 | 7,643 | 0.670 |
| Body Mass | 0.014 | 0.022 | -0.03 | — | 0.058 | 7,582 | 0.515 |
| Daily Survival Rate | 0.006 | 0.022 | -0.038 | — | 0.049 | 7,674 | 0.788 |
| Ambient Temperature x Body Mass | 0.014 | 0.019 | -0.024 | — | 0.052 | 7,127 | 0.476 |
| *Random Effects* on Nest ID | |  |  |  |  |  |  |
| σ Intercept | 0.042 | 0.007 | 0.031 | — | 0.057 |  |  |
| σ Day | 0.027 | 0.004 | 0.02 | — | -0.001 |  |  |
| ρ (Intercept, Day) | -0.008 | 0.004 | -0.016 | — | 0.036 |  |  |

**Table S4. Accounting for phylogenetic relatedness in principal component analysis reveals identical drivers of incubation behavior: temperature, modulated by body mass and daily survival rates.** Phylogenetic generalized linear mixed models of incubation fragmentation (constructed using hpPCA). Daily survival rates were estimated for each nest in the prior logistic exposure models. Models converged with 3 chains, 100,000 samples, 20,000 burn-in, and thinning rate of 30. Within the G-structure, phylogeny has a posterior mean of 1.141, with effective sample size = 8,001. The R-structure (variable effects) has a posterior mean of 0.248, with effective sample size = 8,001. Significant parameters in bold, x denotes interaction.

| Parameter | Posterior Mean | Standard Error | 95% Credible Interval | | | Effective Sample Size | pMCMC |
| --- | --- | --- | --- | --- | --- | --- | --- |
| *Fixed Effects* |  |  |  |  |  |  |  |
| **Ambient Temperature** | -0.785 | 0.138 | -1.054 | — | -0.51 | 8,001 | **<0.001** |
| Body Mass | -0.005 | 0.171 | -0.343 | — | 0.33 | 8,781 | 0.984 |
| **Daily Survival Rate** | 0.258 | 0.091 | 0.075 | — | 0.432 | 8,001 | **0.005** |
| **Ambient Temperature x Body Mass** | -0.259 | 0.127 | -0.509 | — | -0.01 | 8,001 | **0.041** |
| *Random Effects* on Nest ID | |  |  |  |  |  |  |
| σ Intercept | 0.822 | 0.23 | 0.426 | — | 1.332 | 8,001 |  |
| σ Day | 0.126 | 0.031 | 0.076 | — | -0.065 | 8,001 |  |
| ρ (Intercept, Day) | -0.196 | 0.077 | -0.365 | — | 0.196 | 8,001 |  |

**Table S5. Accounting for phylogenetic uncertainty in phylogenetic generalized linear mixed models reveals identical drivers of incubation behavior: temperature, modulated by body mass.** Phylogenetic generalized linear mixed models of incubation fragmentation with 100 chains, each run on a different phylogenetic hypothesis (*82,83*). Daily survival rates were estimated for each nest in the prior logistic exposure models. Models converged with 100 chains, 100,000 samples, 20,000 burn-in, and thinning rate of 30. Within the G-structure, phylogeny has a posterior mean of 0.389, with effective sample size = 207,059. The R-structure (variable effects) has a posterior mean of 0.087, with effective sample size = 267,474. Significant parameters in bold, x denotes interaction.

| Parameter | Posterior Mean | Standard Error | 95% Credible Interval | | | Effective Sample Size | pMCMC |
| --- | --- | --- | --- | --- | --- | --- | --- |
| *Fixed Effects* |  |  |  |  |  |  |  |
| **Ambient Temperature** | -0.484 | 0.088 | -0.656 | — | -0.310 | 266,700 | **<0.001** |
| Body Mass | -0.006 | 0.102 | -0.211 | — | 0.194 | 105,473 | 0.958 |
| **Daily Survival Rate** | 0.141 | 0.062 | 0.018 | — | 0.263 | 266,700 | **0.024** |
| **Ambient Temperature x Body Mass** | -0.167 | 0.081 | -0.324 | — | -0.007 | 191,468 | **0.041** |
| *Random Effects* on Nest ID | |  |  |  |  |  |  |
| σ Intercept | 0.336 | 0.086 | 0.193 | — | 0.529 | 262,570 |  |
| σ Day | 0.069 | 0.014 | 0.046 | — | 0.101 | 266,097 |  |
| ρ (Intercept, Day) | -0.078 | 0.030 | -0.145 | — | -0.029 | 266,700 |  |

**Table S6. Incubation recess duration, ambient temperature and cumulative egg mass drive temperature loss in unattended nests.** Descriptive, interceptless generalized additive model of nest temperature loss of 5,476 incubation recesses over 256 days from 92 nests of 33 species of hummingbirds. Df denotes degrees of freedom, s() denotes nonlinear effect with natural splines, and x denotes interaction. Temperature denotes average ambient temperature during individual incubation recess. Clutch mass denotes total egg mass of eggs in each nest. Degrees of freedom for nonlinear effect = 3.00. Significant terms in bold.

| Parameter | Estimate | Df | Sum of squares | F-value | P-value |
| --- | --- | --- | --- | --- | --- |
| **s(Recess Duration)** | -0.781 | 1 | 56,482 | 23797.884 | **<0.0001** |
| **Temperature** | -0.111 | 1 | 3,513 | 1480.018 | **<0.0001** |
| **s(Recess Duration) x Temperature** | 0.033 | 1 | 1,504 | 633.784 | **<0.0001** |
| s(Recess Duration) x Clutch Mass | 0.260 | 1 | 2 | 0.764 | 0.382 |
| **Temperature x Clutch Mass** | 0.015 | 1 | 19 | 8.174 | 0.004 |
| **s(Recess Duration) x Clutch Mass x Temperature** | -0.015 | 1 | 194 | 81.902 | **<0.0001** |
| Residuals |  | 5,737 | 13,616 |  |  |

**Table S7. Incubation recess duration, ambient temperature and cumulative egg mass drive temperature loss in unattended nests with phylogenetic models.** Confirmatory, interceptless phylogenetic generalized additive mixed model of nest temperature loss of 5,476 incubation recesses over 256 days from 92 nests of 33 species of hummingbirds. Df denotes degrees of freedom, x denotes interaction. Temperature denotes average ambient temperature during individual incubation recess. Clutch mass denotes total egg mass of eggs in each nest. Models converged with 3 chains, 100,000 samples, 20,000 burn-in, and thinning rate of 30. Within the G-structure, phylogeny has a posterior mean of 2.856, with effective sample size = 8,001 and the random effect of nest day has a posterior mean of 1.426, with effective sample size = 8,001. The R-structure (variable effects) has a posterior mean of 0.975, with effective sample size = 8,001. Significant parameters in bold. The spline of basis 2 had the highest positive loading on the nonlinear effect of Recess Duration and was therefore chosen to interact with the other linear terms. Temperature denotes average ambient temperature during individual incubation recess. Clutch mass denotes total egg mass of eggs in each nest. Significant terms in bold.

| Parameter | Posterior mean | 95% Credible Interval | Effective sample size | pMCMC |
| --- | --- | --- | --- | --- |
| **Recess Duration – Spline 1** | -3.778 | -3.923 – -3.364 | 8,001 | **<0.0001** |
| **Recess Duration – Spline 2** | -10.436 | -12.991 – -7.995 | 8,745 | **<0.0001** |
| **Recess Duration – Spline 3** | -3.923 | -4.112 – -3.731 | 7,458 | **<0.0001** |
| **Temperature** | 0.125 | 0.060 – 0.184 | 8,001 | **<0.0001** |
| Recess Duration – Spline 2 x Temperature | 0.105 | -0.046 – 0.249 | 8,793 | 0.163 |
| Recess Duration – Spline 2 x Clutch Mass | 1.653 | -0.121 – 3.535 | 8,870 | 0.077 |
| Temperature x Clutch Mass | 0.002 | -0.042 – 0.045 | 8,584 | 0.949 |
| Recess Duration – Spline 2 x Temperature x Clutch Mass | -0.050 | -0.159 – 0.055 | 8,931 | 0.368 |

**Table S8. Body mass alone influences hummingbird egg mass.** Phylogenetic generalized linear models of egg mass. Significant parameters in bold. Nest height was evaluated first. ⴕ Denotes terms eliminated during model selection. Adjusted R^2^=0.671 and λ = 0.562 in final model.

| Parameter | N | Estimate | Standard Error | T-value | p-value |
| --- | --- | --- | --- | --- | --- |
| Nest Height ⴕ | 84 | 0.005 | 0.024 | 0.193 | 0.847 |
| Temperature ⴕ | 104 | -0.025 | 0.020 | -1.212 | 0.217 |
| Precipitation ⴕ | 104 | -0.026 | 0.020 | -1.277 | 0.205 |
| Latitude ⴕ | 104 | -0.005 | 0.023 | -0.213 | 0.832 |
| Habitat: low ⴕ | 105 | 0.082 | 0.115 | 0.715 | 0.476 |
| Habitat: medium ⴕ | 105 | 0.008 | 0.114 | 0.068 | 0.946 |
| Habitat: high ⴕ | 105 | 0.121 | 0.119 | 1.017 | 0.312 |
| Migration ⴕ | 105 | 0.086 | 0.045 | 1.907 | 0.059 |
| Insularity ⴕ | 106 | 0.024 | 0.073 | 0.330 | 0.742 |
| **Body Mass** | 104 | 0.336 | 0.023 | 14.523 | <0.001 |

**Table S9. Marginally significant relationship between hummingbird incubation period and egg mass.** Phylogenetic generalized linear models of egg mass and incubation period, accounting for allometric effects of body mass. Residuals were generated from phylogenetic log-log regressions of incubation period or egg mass on body mass, respectively. N = 89 species.

| Response | Predictor | Estimate | Standard Error | T-value | p-value | λ | Adjusted  R^2^ |
| --- | --- | --- | --- | --- | --- | --- | --- |
| Residual Incubation Period | Residual Egg Mass | 0.112 | 0.060 | 1.878 | 0.064 | <0.001 | 0.028 |
| Incubation Period | Body Mass | 0.007 | 0.024 | 0.289 | 0.773 | <0.001 | 0.154 |
|  | Egg Mass | 0.044 | 0.234 | 1.866 | 0.066 |  |  |
| Egg Mass | Body Mass | 0.336 | 0.025 | 13.364 | <0.001 | 0.555 | 0.703 |
|  | Incubation Period | 0.030 | 0.021 | 1.403 | 0.164 |  |  |

**Table S10. Effects of phylogenetic uncertainty on comparative models of hummingbird incubation period and egg mass.** Final phylogenetic linear models of incubation period and egg mass, refitted on 100 phylogenetic hypotheses. Medians and range are shown across 100 models are shown in parentheses.

|  | Response | Predictors | Estimate | Standard Error | p-value |
| --- | --- | --- | --- | --- | --- |
|  | ln Incubation Period | |  |  |  |
|  |  | Temperature | -0.038 (-0.041 – -0.034) | 0.010 (0.090 – 0.010) | <0.001 (<0.001 – 0.001) |
|  |  | ln Body Mass | 0.043 (0.038 – 0.044) | 0.009 (0.008 – 0.010) | <0.001 (<0.001 – <0.001) |
|  |  | Insularity | 0.097 (0.088 – 0.106) | 0.030 (0.029 – 0.031) | 0.002 (0.001 – 0.004) |
| ln Egg Mass | | |  |  |  |
|  | | ln Body Mass | 0.328 (0.322 – 0.324) | 0.024 (0.023 – 0.024) | <0.001 (<0.001 – <0.001) |
|  | Residual Incubation Period | |  |  |  |
|  | Residual Body Mass | | 0.095 (0.088 – 0.112) | 0.061 (0.060 – 0.062) | 0.124 (0.064 – 0.155) |

**Table S11. Mayfield estimators of daily survival and predation rates across the elevation gradient.** Not estimated for nests at Zygia as none failed.

| **Mayfield Estimator** | **Location** | **Nest**  **N** | **Nests**  **Failed**  **Or Predated** | **Observation**  **Days** | **Daily**  **Rate** | **SE** | **95% Confidence Interval** | | |
| --- | --- | --- | --- | --- | --- | --- | --- | --- | --- |
| Survival | all | 390 | 262 | 6092 | 0.957 | 0.003 | 0.952 | - | 0.962 |
|  | Remedios | 24 | 16 | 468 | 0.966 | 0.008 | 0.949 | - | 0.982 |
|  | Tatamá | 47 | 30 | 918 | 0.967 | 0.006 | 0.956 | - | 0.979 |
|  | La Minga | 9 | 8 | 209 | 0.962 | 0.013 | 0.936 | - | 0.988 |
|  | Zygia | 2 | 0 | 39 | - | - | - |  | - |
|  | Pantiacolla | 130 | 94 | 1800 | 0.948 | 0.005 | 0.937 | - | 0.958 |
|  | Tono | 39 | 28 | 549 | 0.949 | 0.009 | 0.931 | - | 0.967 |
|  | San_Pedro | 58 | 32 | 937 | 0.966 | 0.006 | 0.954 | - | 0.977 |
|  | Wayqecha | 81 | 54 | 1172 | 0.954 | 0.006 | 0.942 | - | 0.966 |
| Predation | all | 390 | 178 | 6092 | 0.029 | 0.002 | 0.025 | - | 0.033 |
|  | Remedios | 24 | 11 | 468 | 0.024 | 0.007 | 0.01 | - | 0.037 |
|  | Tatamá | 47 | 21 | 918 | 0.023 | 0.005 | 0.013 | - | 0.033 |
|  | La Minga | 9 | 6 | 209 | 0.029 | 0.012 | 0.006 | - | 0.051 |
|  | Zygia | 2 | 0 | 39 | - | - | - |  | - |
|  | Pantiacolla | 130 | 74 | 1800 | 0.041 | 0.005 | 0.032 | - | 0.05 |
|  | Tono | 39 | 19 | 549 | 0.035 | 0.008 | 0.019 | - | 0.05 |
|  | San Pedro | 58 | 18 | 937 | 0.019 | 0.004 | 0.01 | - | 0.028 |
|  | Wayqecha | 81 | 29 | 1172 | 0.025 | 0.005 | 0.016 | - | 0.034 |

**Table S12. Bayesian phylogenetic logistic exposure model to estimate daily nest survival.** Model included data from 6,092 days of observation from 390 nests of 34 species of hummingbirds in which 262 nests failed. Models converged with 3 chains, 4,000 samples, 2,000 burn-in and *adapt_delta* set to 0.95. In random effect terms, variance is denoted by σ and covariance is denoted by ρ. Ȓ denotes Gelman-Rubin statistics.

| Parameter` | Posterior mean | Estimated Error | 95% Credible Interval | | | Ȓ | Bulk Effective Sample Size |
| --- | --- | --- | --- | --- | --- | --- | --- |
| *Fixed Effects* |  |  |  |  |  |  |  |
| Intercept | -3.238 | 0.654 | -4.545 | — | -1.934 | 1.001 | 1219 |
| Elevation | 0.032 | 0.205 | -0.368 | — | 0.438 | 1.001 | 4015 |
| Nest Height | 0.069 | 0.423 | -0.793 | — | 0.871 | 1.005 | 2415 |
| Year: 2008 | 0.976 | 0.711 | -0.418 | — | 2.372 | 1.000 | 1520 |
| Year: 2009 | -0.56 | 0.671 | -1.878 | — | 0.741 | 1.000 | 1423 |
| Year: 2010 | -0.027 | 0.657 | -1.329 | — | 1.248 | 1.000 | 1341 |
| Year: 2011 | -0.297 | 0.685 | -1.659 | — | 1.062 | 1.000 | 1436 |
| Year: 2012 | -0.274 | 0.64 | -1.525 | — | 0.956 | 1.000 | 1316 |
| Year: 2013 | 0.082 | 0.639 | -1.155 | — | 1.34 | 1.000 | 1223 |
| Year: 2014 | -0.095 | 0.647 | -1.326 | — | 1.181 | 1.000 | 1270 |
| Year: 2015 | 0.048 | 0.719 | -1.341 | — | 1.486 | 1.000 | 1493 |
| *Random Effects* | |  |  |  |  |  |  |
| σ Intercept | 0.666 | 0.29 | 0.192 | — | 1.33 | 1.002 | 1955 |
| σ Elevation | 0.755 | 0.397 | 0.164 | — | 1.704 | 1.001 | 2280 |
| ρ (Intercept, Nest Height) | 0.684 | 0.319 | -0.179 | — | 0.994 | 1.001 | 2724 |

**Table S13. Bayesian phylogenetic logistic exposure model to estimate daily predation rate.** Model included data from 6,092 days of observation from 390 nests of 34 species of hummingbirds, in which 178 nests were depredated. Models converged with 3 chains, 4,000 samples, 2,000 burn-in and *adapt_delta* set to 0.95. In random effect terms, variance is denoted by σ and covariance is denoted by ρ. Ȓ denotes Gelman-Rubin statistic.

| Parameter` | Posterior mean | Estimated Error | 95% Credible Interval | | | Ȓ | Bulk Effective Sample Size |
| --- | --- | --- | --- | --- | --- | --- | --- |
| *Fixed Effects* |  |  |  |  |  |  |  |
| Intercept | -1.643 | 0.69 | -2.967 | — | -0.262 | 1.001 | 1575 |
| Elevation | 0.227 | 0.213 | -0.168 | — | 0.658 | 1.001 | 3768 |
| Nest Height | 0.385 | 0.361 | -0.329 | — | 1.121 | 1.002 | 3118 |
| Year: 2008 | 0.236 | 0.767 | -1.241 | — | 1.734 | 1.000 | 2255 |
| Year: 2009 | -1.438 | 0.707 | -2.829 | — | -0.051 | 1.000 | 1807 |
| Year: 2010 | -0.432 | 0.685 | -1.802 | — | 0.899 | 1.000 | 1683 |
| Year: 2011 | -0.886 | 0.699 | -2.277 | — | 0.48 | 1.001 | 1785 |
| Year: 2012 | -0.906 | 0.67 | -2.221 | — | 0.38 | 1.001 | 1636 |
| Year: 2013 | -0.094 | 0.667 | -1.394 | — | 1.193 | 1.001 | 1672 |
| Year: 2014 | -0.208 | 0.667 | -1.512 | — | 1.097 | 1.001 | 1600 |
| Year: 2015 | -0.812 | 0.737 | -2.24 | — | 0.619 | 1.001 | 1772 |
| *Random Effects* | |  |  |  |  |  |  |
| σ Intercept | 0.721 | 0.286 | 0.235 | — | 1.36 | 1.000 | 1740 |
| σ Elevation | 0.654 | 0.337 | 0.11 | — | 1.435 | 1.001 | 2475 |
| ρ (Intercept, Nest Height) | 0.709 | 0.296 | -0.097 | — | 0.993 | 1.000 | 3382 |

**Table S14. Temperature, body mass and daily predation rates influence hummingbird incubation behavior.** Phylogenetic generalized linear mixed models of incubation fragmentation. Daily predation rates were estimated for each nest in the prior logistic exposure models. Models converged with 3 chains, 100,000 samples, 20,000 burn-in, and thinning rate of 30. Within the G-structure, phylogeny has a posterior mean of 0.381, with effective sample size = 9,383. The R-structure (variable effects) has a posterior mean of 0.087, with effective sample size = 8,736. Significant parameters in bold, x denotes interaction.

| Parameter | Posterior Mean | Standard Error | 95% Credible Interval | | | Effective Sample Size | pMCMC |
| --- | --- | --- | --- | --- | --- | --- | --- |
| *Fixed Effects* |  |  |  |  |  |  |  |
| **Ambient Temperature** | -0.485 | 0.086 | -0.654 | — | -0.318 | 8,001 | **<0.001** |
| Body Mass | 0.003 | 0.104 | -0.207 | — | 0.204 | 7,992 | 0.963 |
| **Daily Predation Rate** | -0.163 | 0.064 | -0.289 | — | -0.037 | 8,001 | **0.010** |
| **Ambient Temperature x Body Mass** | -0.172 | 0.081 | -0.329 | — | -0.016 | 9,314 | **0.031** |
| *Random Effects* on Nest ID | |  |  |  |  |  |  |
| σ Intercept | 0.328 | 0.084 | 0.187 | — | 0.518 | 8,001 |  |
| σ Day | 0.069 | 0.014 | 0.046 | — | 0.101 | 7,736 |  |
| ρ (Intercept, Day) | -0.078 | 0.030 | -0.146 | — | -0.029 | 8,001 |  |

**Table S15. Survival does not influence hummingbird incubation period.** Phylogenetic generalized linear models of incubation period. Significant parameters in bold. Daily Survival Rate was evaluated first and Nest Height was evaluated second due to sample size. ⴕ Denotes terms eliminated during model selection. Adjusted R^2^=0.241 and λ = 0.170 in final model.

| Parameter | N | Estimate | Standard Error | T-value | p-value |
| --- | --- | --- | --- | --- | --- |
| Daily Survival Rate ⴕ | 30 | 0.009 | 0.029 | 0.327 | 0.750 |
| Nest Height ⴕ | 128 | 0.008 | 0.011 | 0.786 | 0.434 |
| Precipitation ⴕ | 162 | 0.001 | 0.010 | 0.126 | 0.900 |
| Latitude ⴕ | 162 | -0.008 | 0.009 | -0.948 | 0.345 |
| Habitat: low ⴕ | 162 | -0.019 | 0.040 | -0.467 | 0.641 |
| Habitat: medium ⴕ | 162 | -0.051 | 0.038 | -1.355 | 0.178 |
| Habitat: high ⴕ | 162 | -0.034 | 0.040 | -0.858 | 0.393 |
| Migration ⴕ | 162 | -0.012 | 0.023 | -0.517 | 0.606 |
| **Temperature** | 162 | -0.039 | 0.010 | -4.057 | <0.001 |
| **Body Mass** | 162 | 0.044 | 0.009 | 4.778 | <0.001 |
| **Insularity** | 164 | 0.094 | 0.030 | 3.095 | 0.002 |

**Table S16. Bayesian phylogenetic regression model with measurement error on daily survival rate shows no effect of daily survival rate.** We refitted the starting phylogenetic regression model to include phylogeny as well as measurement error on the term of daily survival rate. Model included data on 21 species of hummingbirds. Models converged with 3 chains, 600,000 samples, 100,000 burn-in, and thinning rate of 500 and *adapt_delta* set to 0.95. In random effect terms, variance is denoted by σ and Ȓ denotes Gelman-Rubin statistics. Measurement error is on Daily Survival Rate is denoted by “DSR|SE”. Daily survival rate has no effect, thereby justifying its exclusion and expansion of the dataset.

| Parameter | Posterior mean | Estimated Error | 95% Credible Interval | Ȓ | Bulk Effective Sample Size |
| --- | --- | --- | --- | --- | --- |
| *Fixed Effects* | |  |  |  |  |
| Body Mass | 0.023 | 0.036 | -0.049 – 0.094 | 1.001 | 2,843 |
| Ambient Temperature | -0.029 | 0.045 | -0.117 – 0.056 | 1.001 | 2,753 |
| Precipitation | -0.027 | 0.058 | -0.146 – 0.086 | 1.001 | 2,947 |
| Latitude | 0.032 | 0.045 | -0.062 – 0.122 | 1.000 | 2,796 |
| Minimum Nest Height | 0.055 | 0.061 | -0.062 – 0.176 | 1.001 | 2,762 |
| Habitat: medium | 0.053 | 0.120 | -0.190 – 0.299 | 1.001 | 2,949 |
| Habitat: high | 0.057 | 0.142 | -0.221 – 0.338 | 1.000 | 2,973 |
| Migration | -0.079 | 0.125 | -0.323 – 0.168 | 1.000 | 2,872 |
| Insularity | -0.052 | 0.203 | -0.435 – 0.348 | 1.001 | 2,925 |
| Daily Survival Rate\|SE | 0.014 | 0.047 | -0.076 – 0.113 | 1.001 | 2,877 |
| Residual σ | 0.068 | 0.046 | 0.006 – 0.175 | 0.999 | 2,856 |
| *Random Effects* | |  |  |  |  |
| σ Intercept | 0.083 | 0.058 | 0.004 – 0.214 | 1.001 | 2,915 |

**Table S17. Bayesian phylogenetic generalized mixed model shows no variation in adult body temperature across elevations.** Using data from (*58*), we explored if adult mean nocturnal internal body temperature varies across an elevational gradient between 520m and 2,500m asl in 112 samples of 24 species of hummingbirds. Models converged with 3 chains, 600,000 samples, 200,000 burn-in, and thinning rate of 1000 and *adapt_delta* set to 0.95. In random effect terms, variance is denoted by σ and Ȓ denotes Gelman-Rubin statistics.

| Parameter | Posterior mean | Estimated Error | 95% Credible Interval | Ȓ | Bulk Effective Sample Size |
| --- | --- | --- | --- | --- | --- |
| *Fixed Effects* | |  |  |  |  |
| Elevation | 0.213 | 0.336 | -0.42 - 0.907 | 1.001 | 1,173 |
| Residual σ | 1.904 | 0.158 | 1.619-2.227 | 1.001 | 1,140 |
| *Random Effects* | |  |  |  |  |
| σ Intercept | 0.671 | 0.515 | 0.028-1.916 | 1.000 | 1,121 |
